# Acute Food Deprivation Rapidly Modifies Valence-Coding Microcircuits in the Amygdala

**DOI:** 10.1101/285189

**Authors:** Gwendolyn G Calhoon, Amy K Sutton, Chia-Jung Chang, Avraham M Libster, Gordon F Glober, Clémentine L Lévêque, G David Murphy, Praneeth Namburi, Christopher A Leppla, Cody A Siciliano, Craig P Wildes, Eyal Y Kimchi, Anna Beyeler, Kay M Tye

## Abstract

In the quest for food, we may expend effort and increase our vulnerability to potential threats. Motivation to seek food is dynamic, varying with homeostatic need. What mechanisms underlie these changes? Basolateral amygdala neurons projecting to the nucleus accumbens (BLA→NAc) preferentially encode positive valence, whereas those targeting the centromedial amygdala (BLA→CeM) preferentially encode negative valence. Longitudinal *in vivo* two-photon calcium imaging revealed that BLA→NAc neurons were more active, while BLA→CeM neurons were less active, following just 1 day of food deprivation. Photostimulating BLA→CeM neurons inhibited BLA→NAc neurons at baseline, but food deprivation rapidly converted this inhibition into facilitation, supporting a model wherein BLA→NAc excitability mediates invigorated food-seeking behavior after deprivation. Indeed, inhibiting BLA→NAc reduced motivation for a caloric reinforcer in food deprived animals. Taken together, negative valence overrides positive valence processing in satiety, but changing homeostatic needs alter reward value via a rapid shift in the balance between projection-defined populations of BLA neurons.

## Introduction

For an animal to engage in reward-seeking behaviors versus defensive behaviors, it must recognize environmental stimuli as potentially rewarding or threatening. The rapid parsing of external cues into “good” or “bad” is aided by amygdala circuits which encode the emotional valence of these environmental cues. However, the interpretation of such cues by the animal as positive or negative depends not only upon some objective quality of the cue or what it predicts, but also upon the individual needs of the animal when it is making the interpretation. A classic example is that a concentrated sodium solution that would normally be aversive is treated as palatable by salt-depleted rats (Berridge et al., 1984; Wagman, 1963). Moreover, if an animal is especially hungry, a cue that predicts a caloric reward might be expected to register more robustly than if the animal is sated (Aitken et al., 2016). Certainly, attention and response bias toward food-associated cues are enhanced by hunger (Burgess et al., 2017). Here, we investigate how a change in internal satiety state impacts amygdala neurons that encode the emotional valence of external stimuli.

The basolateral amygdala (BLA) is a nucleus poised to decipher the predictive value of environmental events and to integrate that information with signals indicating internal state (Burgess et al., 2017; Janak and Tye, 2015; Meyer et al., 2014). Within the BLA, individual populations of neurons encode cues predicting aversive or appetitive outcomes, as well as the emotional valence of stimuli (Gore et al., 2015; Kim et al., 2016; Lee et al., 2017; Namburi et al., 2016; Paton et al., 2006; Quirk et al., 1995; Redondo et al., 2014; Shabel and Janak, 2009; Tye et al., 2008). Multiple lines of evidence support a model wherein two projection-target defined populations of BLA neurons predominantly encode positive and negative valence in an opposing manner (Beyeler et al., 2016; Namburi et al., 2015). First, synaptic input onto BLA→NAc neurons is strengthened following reward learning and weakened after fear conditioning, whereas the opposite changes occur in synapses onto BLA→CeM cells (Namburi et al., 2015). Second, *in vivo* electrophysiological recordings in photoidentified BLA→NAc neurons reveal that they primarily encode positive valence, while BLA→CeM neurons largely encode negative valence (Beyeler et al., 2016). Third, photostimulation of the BLA→NAc projection supports positive reinforcement (Britt et al., 2012; Namburi et al., 2015; Stuber et al., 2011), whereas activating BLA→CeM neurons produces avoidance (Namburi et al., 2015).

In addition to these valence coding functions, BLA neurons have multiple properties consistent with a role in feeding behavior. A number of major input sources to the BLA are critically involved in feeding related signals and have the potential to modulate BLA neural activity in a satiety-dependent manner. For example, the paraventricular thalamus (PVT) densely innervates the BLA and has been shown to contribute to food intake and appetitive motivation (Betley et al., 2013; Do-Monte et al., 2017; Livneh et al., 2017; Millan et al., 2017; Vertes and Hoover, 2008). What is more, the BLA itself has been shown to be sensitive to satiety and food intake; for example, gene expression and activity in the BLA are influenced by palatable food consumption (Packard et al., 2017), and the BLA exhibits local plasticity and structural changes in models of anorexia (Wable et al., 2014). The BLA plays an important role in satiety-dependent outcome devaluation (Corbit et al., 2013; Johnson et al., 2009; Parkes and Balleine, 2013; Shiflett and Balleine, 2010), and mediates hunger-induced invigorated responding (Malvaez et al., 2015). Moreover, the BLA expresses receptors for hunger signals such as ghrelin (Huang et al., 2016; Meyer et al., 2014) and melanocortins (Mountjoy et al., 1994), and sends feedback to sensory regions modulated by satiety state (Burgess et al., 2016; Livneh et al., 2017). Considered together, these data suggest that the BLA is well-positioned to leverage internal information about homeostatic need to guide reevaluation of reinforcers across dynamic conditions and translate value representations into motivated behavior.

Moreover, the projection-target defined populations of BLA neurons that oppositely encode valence are particularly likely to be modulated by satiety state related signals, as the target nuclei of these populations are important for feeding behaviors. The central amygdala (CeA) has been repeatedly demonstrated to play a role in feeding and palatable reward seeking behavior (Carter et al., 2013; Douglass et al., 2017; Han et al., 2017; Holland and Gallagher, 2003; Kim et al., 2017; Robinson et al., 2014; Seo et al., 2016), and the role of NAc in regulating calorically dense food intake is well established (Aitken et al., 2016; Baldo and Kelley, 2007; Counotte et al., 2014; Mietlicki-Baase et al., 2014; O’Connor et al., 2015; Roitman et al., 2004, 2005).

Despite this body of literature, a number of questions remain open. What is the circuit mechanism by which food deprivation increases food-seeking motivation? Would changes in value be determined locally in the BLA or simply relayed through it? If represented locally, how are these changing states dynamically represented in individual neurons? What is the time course of hunger-related modifications to BLA functionality? How do changes in satiety remodel local microcircuitry? Are changes in BLA→ NAc activity necessary for mediating deprivation-induced increases in food-seeking motivation?

Here, we answer these questions using a multifaceted approach. We longitudinally track activity among BLA→ NAc and BLA→CeM neurons across different satiety states *in vivo* using deep brain two-photon calcium imaging. We go on to investigate the interactions among these populations using *ex vivo* slice electrophysiology, and unveil the impact of food deprivation upon these interactions and upon global inputs received by these cells. Finally, we evaluate the necessity of activity in the BLA→ NAc population for motivated seeking of food.

## Results

### Food deprivation rapidly increases basal activity of BLA→NAc neurons and decreases activity in BLA→CeM cells

In order to determine how changes in satiety impact valence-encoding BLA neurons, we tracked calcium activity in individual projection-defined cells across multiple days and different satiety states using endoscopic two-photon imaging. We restricted GCaMP6m expression to BLA→NAc neurons (or BLA→ CeM neurons in a separate cohort of mice) using a dual viral recombination approach, in which we injected a canine adenovirus carrying Cre recombinase (CAV2-Cre) into the projection target structure (NAc or CeM), and infused a virus carrying a Cre-dependent GCaMP6m into the BLA. We then implanted a GRIN lens above the BLA to enable subsequent two-photon imaging of GCaMP6m-expressing cells (Sato et al., 2017; **Figure 1A and E**).

**Figure 1.**
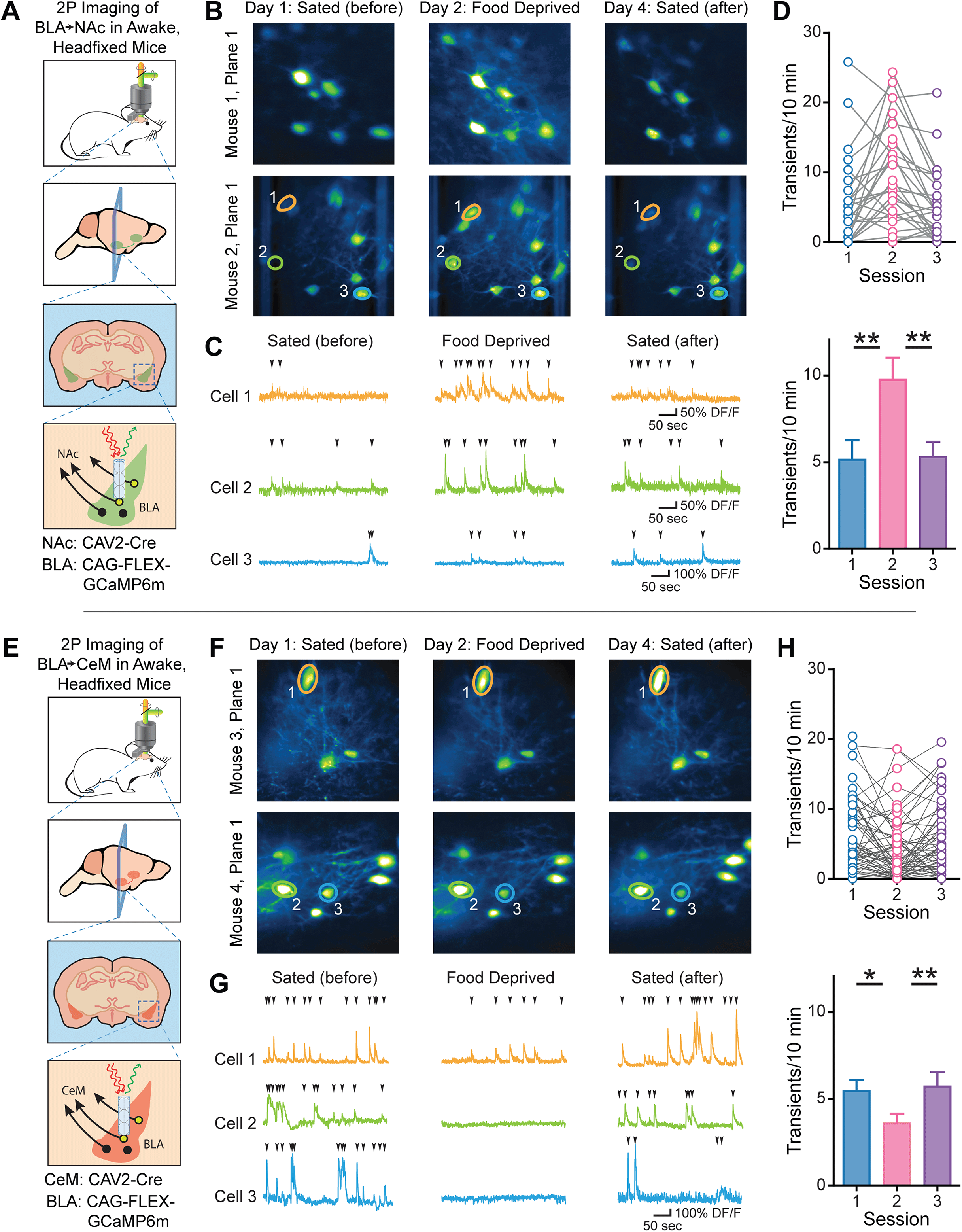
Food deprivation increases BLA→NAc, but decreases BLA→CeM basal activity as observed by longitudinal two-photon imaging. (A) Experimental schematic; we selectively expressed GCaMP6m in BLA→NAc projection neurons and implanted a gradient index (GRIN) lens above the BLA to enable two photon imaging of these cells across multiple days in awake, head-fixed mice. (B) Average fluorescence from two example imaging planes recorded in different individual mice (rows), which were imaged on three separate days (columns). Mice were sated during the first imaging session, and then food deprived for 20 hours leading up to the second imaging session. They were then given free access to food for two days and re-imaged for an additional sated session. (C) GCaMP6m activity in three example neurons from Mouse 2, Plane 1 across the three imaging sessions. The ROIs used to generate these traces are indicated in (B), and carets above each trace indicate events counted as transients. (D) The frequency of calcium transients among BLA→ NAc neurons is sensitive to food deprivation (non-parametric Friedman test of differences among repeated measures, Chi-square=15.50, \*\*\**p*=0.0004), such that more transients occur during session two compared to session one (Dunn’s multiple comparisons test; Rank sum diff.=-27.0, \*\**p*<0.01) and session three (Rank sum diff.=24.0, \*\**p*<0.01). (E) In a separate set of animals, we selectively expressed GCaMP6m in BLA→CeM projection neurons and performed two-photon imaging of these cells through a GRIN lens implanted above the BLA. (F) Example imaging planes from two individual mice that were imaged first while sated, then while food deprived, and finally following 48 hours of free access to food. (G) GCaMP6m activity in three example neurons from the two imaging planes shown above across the three sessions. The ROIs used to generate these traces are indicated in (F), and carets above each trace indicate events counted as transients. (I) The frequency of calcium transients among BLA→CeM neurons is also sensitive to food deprivation (non-parametric Friedman test of differences among repeated measures, Chi-square=11.67, \*\**p*=0.0029), such that fewer transients occur during session two compared to session one (Dunn’s multiple comparisons test; Rank sum diff.=29.0, \**p*<0.05) and session three (Rank sum diff.=-34.0, \*\**p*<0.01). All bar graphs represent the mean, with error bars illustrating SEM. Panels A-D, n=2 mice; panels E-H, n=2 mice.

To test the hypothesis that individual BLA neurons track satiety in a dynamic and reversible manner, we imaged the same projection-defined neurons across sated and hungry conditions. Upon food deprivation, the number of calcium transients increased in BLA→ NAc neurons, and returned to baseline after mice were re-fed (Friedman Test, Chi-square=15.50, \*\*\**p*=0.0004; **Figure 1B-D**). In contrast, we found a decrease in the frequency of calcium transients among BLA→CeM neurons in food deprived mice (Chi-square=11.67, \*\**p*=0.0029; **Figure 1F-H**). As we observed among the BLA→ NAc population, the basal frequency of calcium transients was restored in BLA→CeM neurons within 48 hours by providing ad libitum food access (**Figure 1F-H**).

These findings imply that there may be competition between these functionally-opposed valence-encoding populations. While valence-encoding properties for projection-defined populations of BLA neurons are well-described (Beyeler et al., 2016, 2018), it is yet unknown whether interactions between them inform action selection - or if synaptic contacts even exist. While photoactivation of either of these BLA populations inhibits neighboring BLA neurons *in vivo* (Beyeler et al., 2018), the identity of the inhibited neurons is unknown. Considering that these two populations of neurons are intermingled within the BLA (Beyeler et al., 2018), the opposing changes in calcium activity following food deprivation raise the question of whether the BLA→ NAc and BLA→CeM neurons locally interact.

### Local interactions between BLA→NAc and BLA→CeM neurons

The food-deprivation induced changes in BLA basal activity *in vivo* may reflect global changes in inputs to the BLA as well as locally-mediated microcircuit modifications. To evaluate what, if any, local microcircuit interactions occur between BLA→NAc and BLA→CeM neurons, we used whole-cell patch-clamp recordings in acute slice preparations containing the BLA. To enable targeted patching of BLA neurons, we injected retrogradely-travelling fluorescent microspheres (retrobeads) into either the NAc or the CeM (**Supplemental Figures 2, 3A&D**). In addition to injecting retrobeads, we expressed ChR2-eYFP in the opposing population using a dual viral recombination approach, in which we injected CAV2-Cre into the NAc or CeM and an adeno-associated virus carrying Cre-dependent ChR2-eYFP into the BLA (**Supplemental Figure 2**). Using this strategy, we recorded from BLA→NAc neurons and optically stimulated BLA→CeM neurons (**Supplemental Figure 3A-C**). Conversely in separate animals, we recorded from BLA→ CeM neurons and optically stimulated BLA→NAc neurons (**Supplemental Figure 3D-F**).

**Figure 2.**
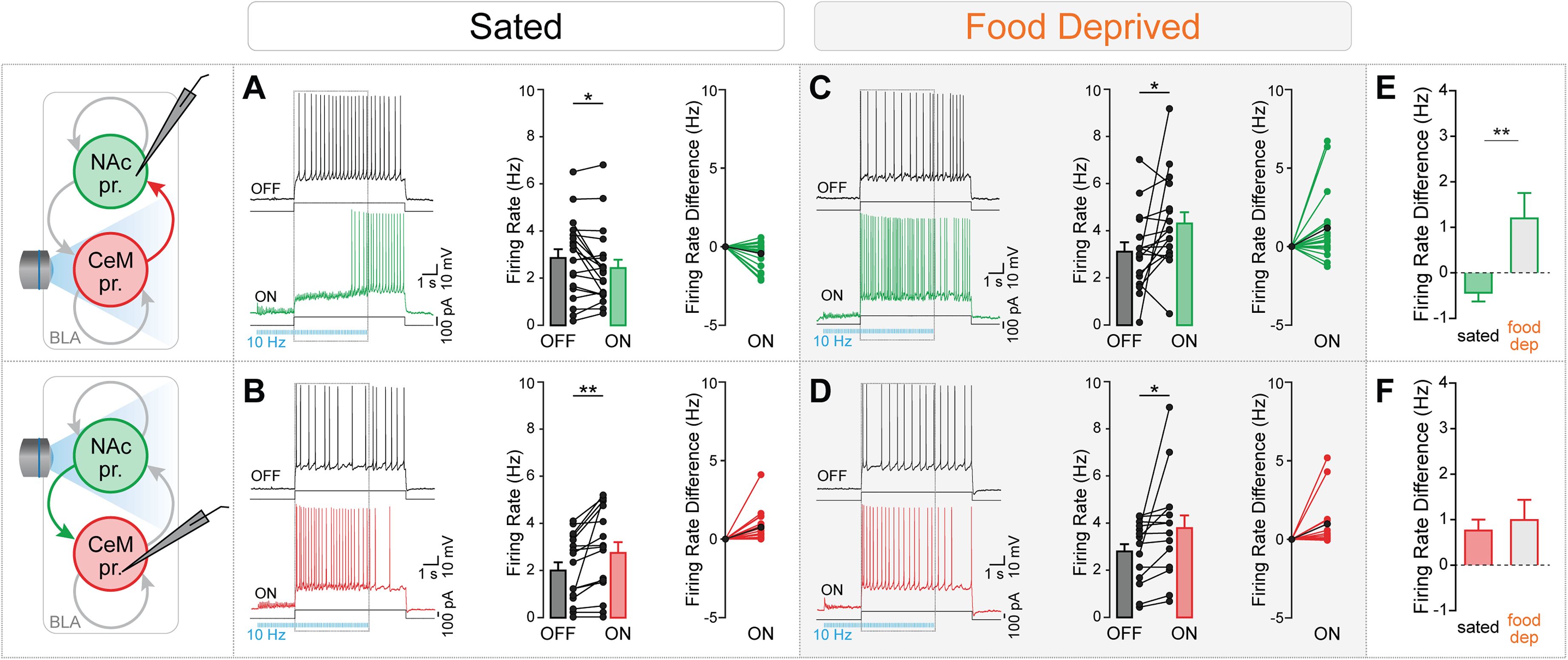
Influence of BLA→CeM neuron photostimulation on BLA→NAc firing is reversed following food deprivation. (A) Photostimulation of BLA→CeM neurons decreases current-evoked firing rate in BLA→NAc neurons in slices taken from sated mice. Current injection through the recording electrode evoked firing in BLA→NAc neurons in the absence of photostimulation (example shown in left panel, black trace (“OFF”); group data shown as black bar in middle panel). Firing rate was decreased by simultaneous 10 Hz photostimulation of BLA→CeM neurons (green trace (“ON”) in left panel, green bar in center panel, green circles in right panel, average change in firing rate indicated by closed black circle in right panel; paired t-test, t_19_=2.238, \**p*=0.0374). n=8 mice, 20 cells. (B) Photostimulation of BLA→NAc neurons increases current-evoked firing in BLA→CeM neurons in slices taken from sated mice (paired t-test, t_16_=3.003, \*\**p*=0.0084). n=7 mice, 17 cells. (C) In slices taken from mice food deprived 20 hours prior to slicing, BLA→CeM stimulation facilitates current-evoked firing in BLA→NAc neurons (paired t-test, t_16_=2.172, \**p*=0.0452). n=6 mice, 17 cells. (D) BLA→NAc photostimulation increases the current-evoked firing rate of BLA→CeM neurons following food deprivation (paired t-test, t_13_=2.241, \**p*=0.0432). n=6 mice, 15 cells. (E) Food deprivation changes the impact of BLA→CeM photostimulation on current-evoked firing in BLA→NAc neurons *(Mann-Whitney U=76.5,* \*\**p*=0.0036), such that there is a shift from an inhibitory to a facilitative relationship following food deprivation. (F) BLA→NAc photostimulation increases current-evoked firing among BLA→CeM neurons regardless of satiety state *(Mann-Whitney U*=116.5, *p*=0.9297). Dotted rectangles in the example traces shown in (A)-(H) represent the time period of blue light delivery during a current injection, which corresponds to the epochs used to compare firing rates. Bar graphs illustrate mean with SEM. In firing rate difference plots, mean change in firing rate is indicated in black circles.

To determine whether functional monosynaptic contacts exist between these populations, we recorded BLA→NAc neurons in voltage-clamp, and in all cases (n=12), 5 ms photostimulation of BLA→CeM neurons resulted in an excitatory post synaptic current (EPSC) when holding the cell at −70 mV and an inhibitory post synaptic current (IPSC) when holding the cell at 0 mV (**Supplemental Figure 3A**). Although the relative amplitude of these two currents did not differ in the cells we sampled (Wilcoxon signed-rank test, *p*=0.3804), the onset latency of the EPSC was shorter than that of the IPSC (t_10_=4.540, \*\**p*=0.0011), suggesting a monosynaptic and polysynaptic response, respectively (**Supplemental Figure 3A**). To determine if the EPSC measured in BLA→ NAc neurons following BLA→CeM photostimulation was indeed monosynaptic, we used ChR2-assisted circuit mapping (Petreanu et al., 2007) and found that the EPSC persisted following bath application of 1μM tetrodotoxin and 100 μM 4-Aminopyridine (TTX/4AP). Additionally, the EPSC was glutamatergic, as indicated by its persistence in the presence of 100 μM picrotoxin (PTX). EPSC amplitude was significantly decreased by 20 μM 2,3-dihydroxy-6-nitro-7-sulfamoyl-benzo[f]quinoxaline-2,3-dione (NBQX; t_7_=2.989, \**p*=0.0202; **Supplemental Figure 3B**), indicating that AMPAR-mediated currents comprised the bulk of the EPSC. In half of the BLA→ NAc neurons we recorded, we evaluated the impact of PTX upon the optically-evoked IPSC, whereas in the other half, we tested the effect of TTX/4AP upon this current. The IPSC was eliminated both by PTX and TTX/4AP (t_6_=2.729, \**p*=0.0342 and t_4_=5.330, \*\**p*=0.0060, respectively, **Supplemental Figure 3C**), suggesting that, in addition to the monosynaptic glutamatergic response evoked in BLA→NAc neurons following photostimulation of the BLA→CeM population, BLA→CeM neurons also recruit an inhibitory intermediary to elicit a GABAergic current in BLA→ NAc neurons.

To examine the impact of optically-evoked activity in the BLA→ NAc population upon BLA→CeM neurons, we recorded retrobead-positive BLA→CeM neurons and stimulated BLA→NAc neurons expressing ChR2-eYFP using 5 ms pulses of blue light (**Supplemental Figure 3D**). In all of the BLA→CeM neurons we recorded in voltage-clamp (n=11), BLA→NAc photostimulation elicited EPSCs at −70 mV and IPSCs at 0 mV, and onset latencies for EPSCs were shorter than those for IPSCs (t_11_=4.289, \*\**p*=0.0016; **Supplemental Figure 3D**). Parallel to the recordings we performed in BLA→NAc neurons, we observed that the EPSC evoked in BLA→CeM neurons was not eliminated by PTX nor TTX/4AP, but bath application of NBQX significantly reduced the magnitude of this current (t_7_=4.814, \*\**p*=0.0019; **Supplemental Figure 3E**). Moreover, the IPSC was sensitive to both PTX and TTX/4AP (t_6_=4.078, \*\**p*=0.0065 and t_3_=2.123, *p*=0.1239, respectively; **Supplemental Figure 3F**). Together, these observations indicate that photoactivation of BLA→NAc neurons elicits a monosynaptic glutamatergic response, as well as a polysynaptic GABAergic response in all BLA→CeM neurons recorded.

Surprisingly, we observed both an EPSC and an IPSC in every cell recorded from BLA→NAc and BLA→CeM cells when photostimulating the opposing population. We were not anticipating that we would observe such dense, widespread innervation of both monosynaptic excitation and feed-forward inhibition onto every cell sampled, based upon the sparse connectivity of BLA neurons reported from paired slice recordings (Duvarci and Pare, 2014). However, it seems likely that our approach of population-level stimulation enhanced our ability to detect interactions among these BLA projection neurons, so we continued to use this strategy moving forward.

Based upon our observations from these voltage-clamp experiments, we concluded that both BLA→NAc and BLA→CeM neurons form synaptic contacts on cells in the opposing population, and that both populations exert inhibition upon each other (likely via local interneurons). However, on their own, these data do not clarify how activity in one projection-target defined population may impinge upon activity in the other, and thereby ultimately shape downstream signaling. Given that we previously observed inhibition of neighboring BLA neurons following photostimulation of these populations *in vivo* (Beyeler et al., 2018), and observed opposing changes in calcium activity in these cells following food deprivation, we predicted that BLA→NAc and BLA→CeM neurons would mutually inhibit each other’s firing. To address this essential question, we recorded a separate set of cells in current-clamp to evaluate how photostimulation of each population modulates action potential firing in the other.

We developed our photostimulation paradigm based upon the firing properties we observed in BLA neurons. We evaluated basal membrane features in all recorded cells, which were consistent with what we have reported previously (Namburi et al., 2015) and largely did not differ between the two populations (**Supplemental Figure 4**). BLA→CeM neurons reached a lower maximal sustained firing rate than BLA→ NAc neurons in response to direct somatic current injection in slices taken from sated animals *(Mann-Whitney U=240.5,* \*\**p*=0.0019; **Supplemental Figure 4C**). Among both populations, however, cells readily maintained sustained firing rates of 10 Hz, so we used trains of 10 Hz photostimulation to elicit robust, but physiologically relevant activity in ChR2-eYFP positive neurons.

To examine how activation of BLA→CeM neurons impacts action potential firing in BLA→NAc neurons, we recorded retrobead positive BLA→NAc cells in BLA slices from mice expressing ChR2-eYFP in BLA→CeM cells (**Figure 2A**). We injected current through the recording electrode in order to evoke firing in the recorded cell, and interleaved sweeps in which we photostimulated the BLA→CeM population using a 7.5 s, 10 Hz train of 5 mW, 5 ms blue LED pulses beginning 2.5 seconds before the onset of the current injection (light ON condition) with sweeps in which the LED controller was turned off (light OFF condition). In this way, we were able to compare the average firing rate during the same portion of the current injection in the presence and absence of BLA→CeM activation. We found that among the BLA→NAc neurons we recorded (n=20), photostimulation of the BLA→CeM population reduced the current-evoked firing rate (t_19_=2.238, \**p*=0.0374; **Figure 2A**). This finding of BLA→CeM-driven inhibition of BLA→NAc neurons is consistent with our observation of robust inhibition of neighboring BLA neurons following optical stimulation of BLA→CeM cells *in vivo* (Beyeler et al., 2018).

It is possible, however, that activation of any subpopulation of BLA neurons would result in the recruitment of local interneurons and subsequently reduce firing in BLA→NAc neurons. To determine whether BLA→NAc neurons are uniquely inhibited by BLA→CeM neurons, we patched BLA→NAc cells from mice in which both the retrobead and CAV2-Cre injections were targeted to the NAc. We avoided conflating within-population interactions among BLA→NAc neurons with direct effects of ChR2 activation in recorded cells by rejecting any cells responding to a one second LED stimulus with photocurrents (data not shown). Photostimulation of BLA→ NAc neurons exerted diverse effects upon the current-evoked firing rate of ChR2-eYFP negative BLA→ NAc neurons, such that the net consequence was no significant change in firing rate (t_15_=2.033, ^+^*p*=0.0601; **Supplemental Figure 5A**). In contrast, within-population photostimulation of BLA→CeM neurons significantly increased current-evoked firing in neighboring ChR2-eYFP negative BLA→CeM neurons (t_16_=3.745, \*\**p*=0.0018; **Supplemental Figure 5B**).

Because the synaptic currents between the two projection-target defined populations were similar, we predicted that modulation of BLA→CeM firing by BLA→ NAc activation would mirror the inhibition of BLA→ NAc neurons following activation of BLA→CeM cells. Surprisingly, we found the opposite to be true. Among the BLA→CeM neurons we recorded (n=17), photostimulation of BLA→NAc neurons increased the current-evoked firing rate (t_16_=3.003, \*\**p*=0.0084; **Figure 2B**).

Importantly, activation of each population modulated firing evoked by current injection in the opposing population; photostimulation of neither population directly elicited spiking in the other. 10 Hz optical stimulation of the BLA→NAc population did not increase firing in BLA→CeM neurons (light only vs basal conditions), nor did optical stimulation of the BLA→CeM population elicit firing in BLA→NAc neurons (light only vs basal conditions) in the absence of a current step (**Supplemental Figure 6**).

### Asymmetric BLA microcircuit interactions depend on satiety state

It is possible that the asymmetric interaction between BLA→NAc and BLA→CeM neurons favor promotion of the BLA→CeM pathway in sated animals because negative valence encoding cells might be necessary for invigorating defensive and escape related behaviors in the face of threat. We hypothesized that the relationship between these two populations could change dynamically depending on the needs of the animal, such that hunger might promote food seeking despite the presence of potential threats. To test whether the local interactions between BLA→NAc and BLA→CeM neurons flexibly change depending upon the satiety state of the animal, we repeated the current clamp patching experiments in slices taken from mice food deprived for 20 hours prior to slicing.

In contrast to what we observed in slices taken from sated mice, we discovered that BLA→ NAc neurons were not inhibited by BLA→CeM neurons following food deprivation, but rather the current evoked firing rate in these cells increased following BLA→CeM photostimulation (t_16_=2.172, \**p*=0.0452; **Figure 2C**). Additionally, whereas within-population stimulation of BLA→NAc neurons had a non-significant impact upon current-evoked firing in neighboring BLA→ NAc neurons in slices taken from sated animals, food deprivation unified the activity of this population such that stimulation of ChR2-positive BLA→ NAc neurons caused a robust increase in the current-evoked firing rate of ChR2-negative BLA→NAc neurons (t_15_=2.217, \**p*=0.0425; **Supplemental Figure 5C**). As was the case in sated animals, BLA→CeM neurons showed elevated current-evoked firing following photostimulation of BLA→NAc neurons in the food deprivation condition (t_13_=2.241, \**p*=0.0432; **Figure 2D**), and cells in this population also evidenced facilitation of firing following within-population stimulation in slices taken from food deprive mice (t_13_=3.251, \*\**p*=0.0063; **Supplemental Figure 5D**).

Comparing the magnitude of the change in firing rate between light OFF and light ON conditions following satiety and food deprivation revealed that a change in satiety did not affect the response of BLA→CeM neurons to either BLA→NAc or within-population stimulation *(Mann-Whitney U*=116.5, *p*=0.9297 and *Mann-Whitney U=78.0, p*=0.1085, respectively; **Figure 2F** and **Supplemental Figure 5F**). Although there was a qualitative change in the impact of within-population stimulation of BLA→NAc neurons, food deprivation did not significantly alter the response of ChR2-negative BLA→ NAc neurons to photostimulation of neighboring ChR2-positive BLA→NAc neurons *(Mann-Whitney U*=115.5, *p*=0.6491; **Supplemental Figure 5E**). Food deprivation dramatically altered the response of BLA→NAc neurons to BLA→CeM neurons, however, insomuch as stimulation of the later population inhibited current evoked firing in the former in the sated condition, but facilitated current evoked firing following food deprivation *(Mann-Whitney U=76.5,* \*\**p*=0.0036; **Figure 2E**).

Whereas we originally predicted that both populations would be equally weighted and potentially mutually inhibit one another (**Supplemental Figure 7A**), we observed that the preferentially negative valence encoding BLA→CeM population supersedes activity in the preferentially positive valence encoding BLA→NAc population in sated animals (**Supplemental Figure 7B**). However, a change in internal energy homeostasis shifts the balance between these BLA neurons such that BLA→NAc neurons are no longer inhibited by the BLA→CeM neurons (**Supplemental Figure 7C**). In fact, the changes that we observed in BLA local interactions centered upon BLA→ NAc cells, in that both the qualitative change in within-population interactions as well as the significant change in opposing population interactions were reflected in an alteration of firing among BLA→NAc neurons.

The impact of food deprivation upon activity in the BLA→NAc population was not easily explained by a change in membrane excitability among these cells. There were no significant differences in the majority of the intrinsic properties that we measured between BLA→NAc and BLA→CeM neurons taken from either sated or food deprived mice (**Supplementary Figure 4**), although there was a trend toward interaction between projection target and satiety state in holding current among these groups (F_1,117_=3.413, ^+^*p*=0.0671; **Supplementary Figure 4A**). Food deprivation did impact the firing-rate responses of BLA projection neurons to increasing current steps; whereas the maximum evoked firing rate differed between BLA→NAc and BLA→CeM neurons recorded in slices taken from sated mice, that difference was abated by food deprivation *(Mann-Whitney U*=262.0, *p*=0.3501; **Supplementary Figure 4C**).

### Food deprivation increases quantal release events onto BLA→NAc neurons

Because the changes in the BLA microcircuit we observed following food deprivation were only partially explained by changes in intrinsic excitability among BLA→NAc neurons, we hypothesized that food deprivation leads to an increase in presynaptic glutamate release onto this population of cells. To test this hypothesis, we recorded miniature excitatory post-synaptic currents (mEPSCs) in BLA→NAc and BLA→CeM neurons in slices taken from sated and food deprived mice. Consistent with the results of our current clamp experiments, we detected no differences in mEPSC frequency or amplitude in BLA→CeM neurons following satiety or food deprivation (amplitude: Mann Whitney U=74.00, *p*=0.1845, frequency: Mann Whitney U =102.5, *p*=0.9235; **Figure 3B, D, & F**). Moreover, although there was no difference in mEPSC amplitude in BLA→NAc neurons recorded from sated and food deprived mice (Mann Whitney U =94.00, *p*=0.4680; **Figure 3A&C**), the frequency of mEPSCs in BLA→NAc cells increased following food deprivation (Mann Whitney U =57.00, \**p*=0.0212; **Figure 3A&E**). These data strongly suggest that food deprivation leads to an increase in presynaptic glutamate release specifically onto BLA→NAc neurons, which could in turn enable these cells to overcome the feed-forward inhibition exerted over them by BLA→CeM activation and shift the balance toward monosynaptic excitation from these afferents. Moreover, an increase in upstream excitation could invigorate the overall increase in activity we observed *in vivo* among BLA→NAc neurons using two-photon imaging.

**Figure 3.**
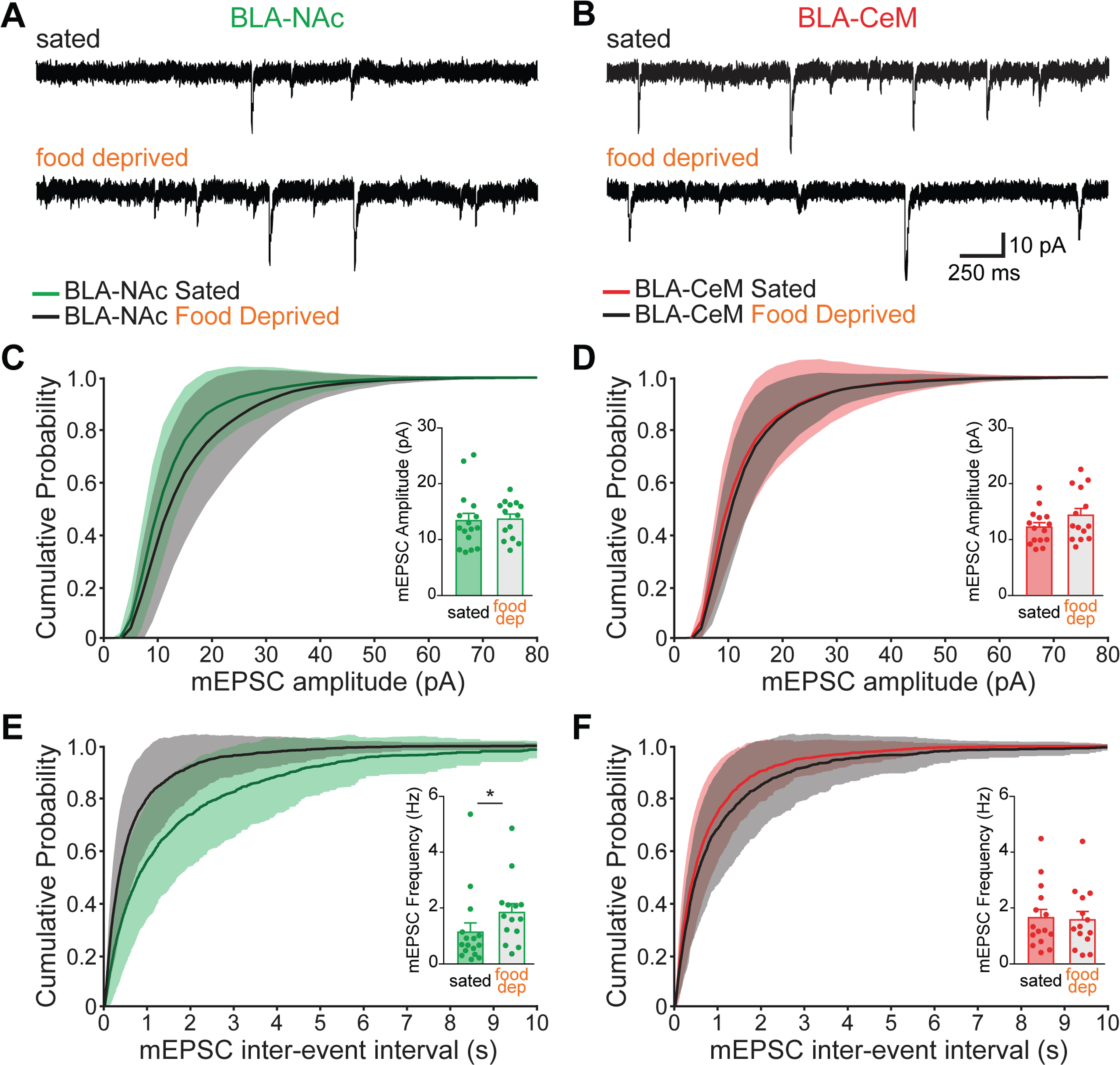
One day of food deprivation increases mEPSC frequency in BLA→NAc, but not BLA→CeM neurons. (A) Example traces containing mEPSCs recorded from BLA→NAc neurons taken from sated (top) and food deprived (bottom) mice. (B) Example mEPSCs recorded from BLA→CeM neurons in slices taken from sated (top) and food deprived (bottom) mice. (C) Cumulative distribution of mEPSC amplitudes in BLA→NAc neurons in slices taken from sated (green) or food deprived (black) mice. Inset: satiety does not affect mEPSC amplitude in BLA→NAc neurons *(Mann-Whitney U*=94.00, *p*=0.4680). Sated: n=4 mice, 16 cells; food deprived: n=3 mice, 14 cells. (D) There is no difference in mEPSC amplitude in BLA→CeM neurons from sated and food deprived mice *(Mann-Whitney U*=74.00, *p*=0.1845). (E) The frequency of mEPSCs is greater in BLA→NAc neurons recorded in slices from mice that had been food deprived than in BLA→NAc neurons from mice that were sated *(Mann-Whitney U=57.00,* \**p*=0.0212). Sated: n=4 mice, 15 cells; food deprived: n=4 mice, 14 cells. (F) Food deprivation did not impact mEPSC frequency in BLA→CeM neurons *(Mann-Whitney U*=102.5, *p*=0.9235). Shaded region represents SEM on cumulative probability plots. All bar graph insets display mean with SEM.

### Increased activity among BLA→NAc neurons is causally linked to increased food seeking behavior

We find that food deprivation rapidly alters the activity and interactions of valence coding BLA neurons, and in particular causes an increase of excitatory input onto the predominantly positive valence encoding BLA→ NAc population. It may be the case that increased activity among BLA→NAc neurons is necessary for invigorating food seeking behavior in hungry mice. In order to test this hypothesis, we employed a chemogenetic approach to inhibit activity in the BLA→NAc population in food deprived mice.

Although the action of Clozapine N-Oxide (CNO) at designer receptors exclusively activated by designer drugs (DREADDs) has recently met with controversy (Gomez et al., 2017), we took several measures to safe-guard the interpretability of our results. First, we include a fluorophore-only control group to enable the detection of any non-specific effects of systemic CNO administration. Second, we use a mid-range dosage of CNO (1 mg/kg) to reduce the probability that efficacious concentrations of metabolites, such as clozapine, accumulate in the animals’ bloodstream. Finally, the timeframe of behavioral measurements that we use is consistent with a range in which CNO has been shown to be a suitable agonist for DREADDs (Mahler and Aston-Jones, 2018). Moreover, we verified the efficacy of CNO to hyperpolarize neurons expressing inhibitory DREADDs (hM4Di) in *ex vivo* whole-cell patch-clamp recordings, and further found that CNO had no impact on the membrane properties of mCherry neurons not expressing hM4Di (**Supplemental Figure 8**).

To achieve the selective expression of the inhibitory DREADD hM4Di bilaterally in BLA→NAc neurons, we used a dual viral recombination approach similar to those described above. Here, we injected CAV2-Cre bilaterally into NAc and infused an adeno associated virus carrying cre-dependent hM4Di-mCherry (or mCherry only in control animals) bilaterally into BLA (**Supplemental Figure 9**). After at least ten weeks to allow for viral expression, we used a within-subjects design to evaluate the necessity of BLA→NAc activity for food intake. Mice were single housed and fasted at the start of their light cycle. Thirty minutes before the start of the dark cycle, we administered either 1 mg/kg CNO or vehicle intraperitoneally. We then provided each mouse with a pre-weighed food pellet at the start of the dark cycle, and weighed the pellet after one and four hours (**Figure 4A**). The mice were re-grouped and returned to *ad libitum* feeding, and we allowed one week between repetitions to ensure full wash-out of CNO and normalization of circadian feeding behaviors. We found that CNO did not significantly impact food intake in hM4Di-expressing mice or mCherry controls within the first hour of feeding (t_4_=1.101, *p*=0.3327 and t_7_=0.0, *p*>0.9999, respectively), nor did CNO impact total food intake in either group (hM4Di: t_4_=0.8513, *p*=0.4426, mCherry: t_7_=0.5298, *p*=0.6126; **Figure 4A**).

**Figure 4.**
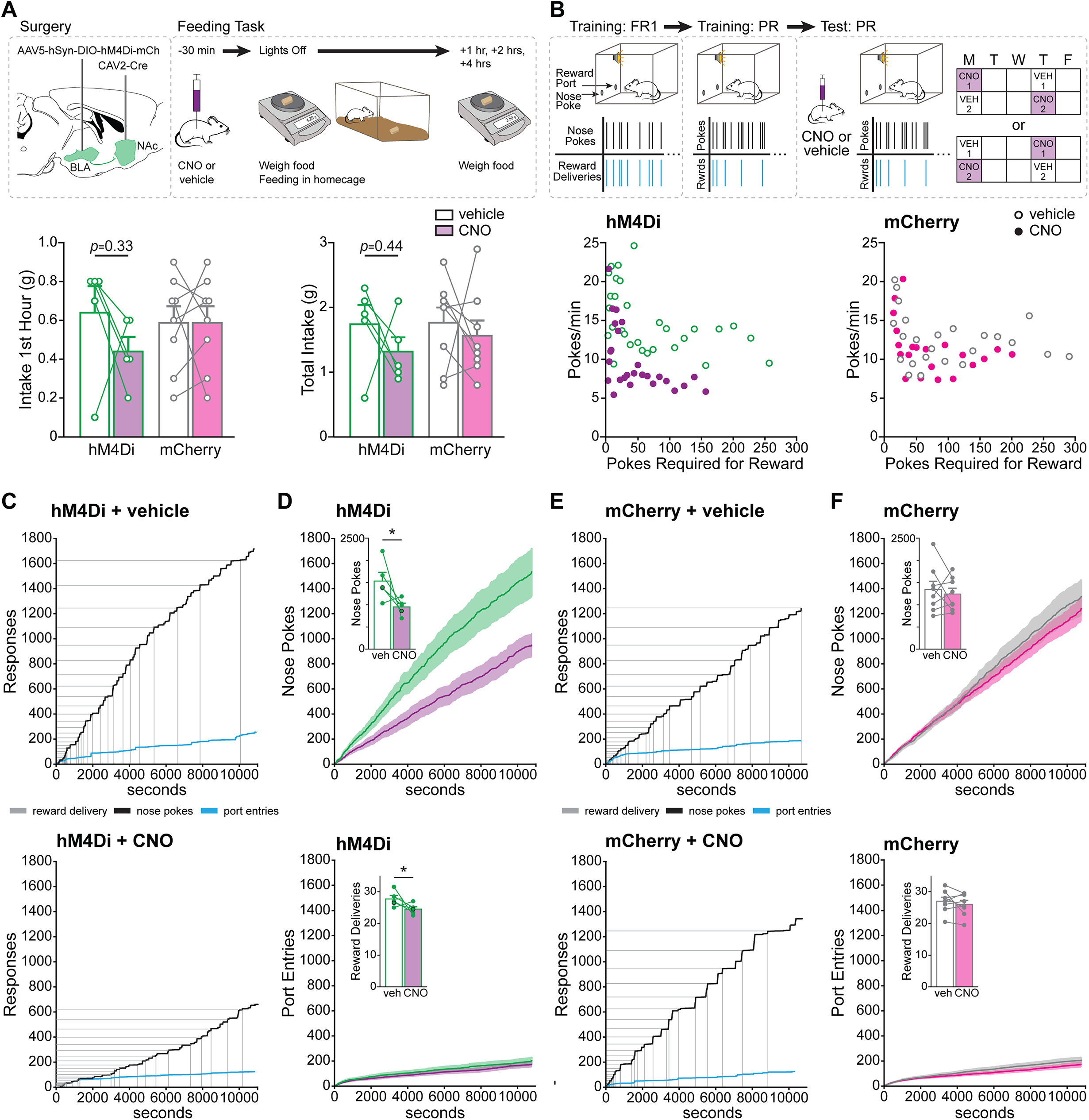
DREADDs inhibition of BLA→NAc neurons reduces appetitive motivation, but not feeding. (A) Top panel: Schematic illustrating the surgical strategy and feeding task paradigm. We selectively expressed inhibitory DREADDs in BLA→ NAc projection neurons bilaterally. We used a within-subjects design for the feeding task, in which the order of CNO or vehicle delivery was counterbalanced. Bottom panel: There was no significant impact of CNO on chow intake among hM4Di-expressing mice (first hour: paired t-test, t_4_=1.101, *p*=0.3327; total intake: paired t-test, t_4_=0.8513, *p*=0.4426) or mCherry controls (first hour: paired t-test, t_7_=0.0, *p*>0.9999; total intake: paired t-test, t_7_=0.5298, *p*=0.6126). (B) Top panel: Schematic representing the approach for progressive ratio training and testing. Mice were first trained on a fixed ratio (FR1) schedule to nose poke for palatable reward delivery, and then graduated to a progressive ratio (PR) schedule. Each mouse completed two vehicle and two CNO progressive ratio sessions, which were counterbalanced. Bottom panel: The rate of responding decreased as the nose pokes required for each subsequent reward increased. This relationship was sensitive to CNO in mice expressing hM4Di in BLA→NAc neurons (left), but not in mCherry controls (right). (C) The performance of an example animal from the hM4Di group on the progressive ratio task following administration of vehicle (top panel) or CNO (bottom panel). Cumulative nose pokes in the operant port are indicated in black, cumulative entries into the reward port are indicated in blue, and reward deliveries are indicated in gray. (D) Top panel: Mean cumulative nose pokes into the operant port for all mice expressing hM4Di in BLA→NAc neurons. Green indicates pokes following vehicle administration; purple indicates pokes following CNO administration. Inset: chemogenetic inhibition of BLA→NAc neurons reduced the mean number of operant nose pokes during the progressive ratio test session (t-test, t_s_=2.7332, \**p*=0.0257). Bottom panel: Mean cumulative entries into the reward port for all mice in the hM4Di group. Inset: the mean number of reward deliveries in a session was reduced following CNO administration relative to vehicle (t-test, t_8_=2.3918, \**p*=0.0437). The example mouse depicted in (C) is indicated with black circles in the insets. (E) Progressive ratio performance of a mouse expressing mCherry in BLA→NAc neurons following vehicle (top panel) or CNO (bottom panel) administration. (F) Mean cumulative nose pokes in the operant port (top panel) and reward port entries (bottom panel) for all mice expressing mCherry in BLA→NAc neurons. Gray: vehicle sessions; pink: CNO sessions. Insets: CNO did not impact the number of nose pokes made by mCherry-expressing mice in the progressive ratio task (top panel; t-test, t_14_=0.4481, *p*=0.6609), nor did it affect the total number of reward deliveries during the task (bottom panel; t-test, t_14_=0.5698, *p*=0.5779). All bar graphs depict mean and SEM. Shaded areas in cumulative plots indicate SEM. In all panels, hM4Di: n=5 mice; mCherry: n=8 mice.

While increased activity among BLA→ NAc neurons does not alter feeding behavior in the homecage, we reasoned that increased activity in a positive-valence encoding population may incite the motivational drive to obtain food reward. We evaluated the necessity of elevated BLA→NAc activity for motivation to obtain a palatable reward using a progressive ratio task, in which the mice were required to expend progressively greater effort to receive each subsequent delivery of a palatable, calorically dense reward (50% Ensure, 50% water solution; **Figure 4B**). Again, we used a within-subjects design, and we repeated both the drug and the vehicle conditions twice to obtain averaged measurements (**Supplemental Figure 10**). For the duration of training, the mice were fed *ad libitum,* but before each test day, we food deprived the mice overnight.

Overall, we found that chemogenetic inhibition of BLA→NAc cells reduced the motivation of hungry mice to obtain a palatable, calorically dense reward (**Figure 4B-F**). Several points of evidence support this conclusion. First, the rate of operant responding in hM4Di-expressing mice markedly decreased with higher reinforcer requirements when the mice were administered CNO (**Figure 4B**). By contrast, the rate of nose poking was not affected by CNO in mCherry-expressing controls. Second, in animals expressing hM4Di in BLA→NAc neurons, CNO administration significantly reduced the total number of operant nose pokes in the progressive ratio session (t_8_=2.7332, \**p*=0.0257; **Figure 4D**). Furthermore, hM4Di-expressing mice earned fewer reinforcer deliveries following CNO administration (t_s_=2.3918, \**p*=0.0437; **Figure 4D**). In mCherry-expressing control mice, CNO administration did not impact the number of operant nose pokes in the session (t_14_=0.4481, *p*=0.6609; **Figure 4F**), or the number of reinforcer deliveries they received (t_14_=0.5698, *p*=0.5779; **Figure 4F**).

Together, these data indicate that while BLA→NAc neurons do not play a direct role in coordinating feeding in food deprived mice, increased activity among these cells is necessary for the appetitive motivation to obtain food rewards.

## Discussion

We provide here the first evidence that the activity of valence encoding populations of BLA neurons targeting the NAc and CeM is impacted by satiety state. Additionally, we reveal the asymmetrical interactions among these populations, and show that these, too, are altered by food deprivation. The bulk of the changes we observed occurred in the predominantly positive valence encoding BLA→NAc population; food deprivation increased calcium activity in these cells, promoted elevated action potential firing in these cells, and increased the rate of presynaptic glutamate release onto these cells. Finally, we demonstrate that inhibiting BLA→NAc neurons decreases motivation of hungry animals to work for a calorically dense food reward.

### State-dependent activity revealed by tracking neurons across days with two-photon imaging

In order to determine whether projection-target defined BLA neurons were sensitive to satiety state, we tracked GCaMP6m activity across multiple days when the animals were fed *ad libitum* or food deprived. There have been notable recent efforts to image amygdala neurons over prolonged periods using single-photon methods (Douglass et al., 2017; Grewe et al., 2017; Li et al., 2017; Yu et al., 2017). However, to track the same individual neurons across multiple sessions over this time period with confidence in our preparation, we concluded that two-photon imaging was required. As the BLA is a deep nucleus, we imaged the cells through a chronically-implanted GRIN lens. This approach enabled us to visually identify individual BLA neurons, which in some cases were silent on one or more sessions and would not have been identifiable using any other approach. This pioneering use of two-photon imaging across multiple days in the BLA revealed that BLA→NAc calcium activity reversibly increases during food deprivation in awake mice, and that by contrast, BLA→CeM calcium activity decreased when the animals were deprived of food.

### Local interactions among projection-defined neurons in the BLA

An important major finding in the current work is that projection-defined valence-encoding BLA populations locally interact to influence each other’s activity. Leading up to this work, it was unknown whether these cells impact each other’s activity, and what the nature of their relationship might be. From an ethological perspective, situations in nature are rarely exclusively rewarding or aversive; for example, animals must often oppose conspecifics and risk injury when seeking mates, or expose themselves to predatory threats when foraging for food or water. This competition between reward and aversion requires interplay between opposing valence systems, potentially at the level of the BLA. Indeed, functionally-defined opposing populations of BLA neurons have been observed to produce differential activity during periods of reward seeking and escape during a naturalistic foraging task in which rats must navigate an environment to obtain food and avoid encountering a mechanical predator (Amir et al., 2015). Based upon the opposing, bidirectional modulation of these populations following fear and reward learning, we hypothesized that BLA→NAc and BLA→CeM neurons mutually inhibit each other, but were surprised by our empirical observations.

Despite the universal existence of both inhibitory and excitatory connections between BLA→NAc and BLA→CeM populations, examination of the net effect of photostimulation of these neurons indicated that they asymmetrically modulate each other’s activity. When we recorded cells in slices taken from sated animals, we found that BLA→CeM neurons, which we have previously characterized as preferentially signaling negative valence, readily suppressed firing in BLA→NAc neurons, which generally signal positive valence. By contrast, BLA→NAc neurons not only failed to inhibit firing in BLA→CeM neurons, they *facilitated* firing in the opposing population. These relationships are schematized in **Supplemental Figure 7**, as are the within-population interactions we observed for both sets of BLA neurons. Considering the individual contributions of these two populations to valence encoding, our present findings provide a potential mechanism allowing fear- and avoidance-related behaviors to supersede reward-seeking in the presence of threat in sated animals.

However, it is maladaptive for animals to exclusively allow threat avoidance to “win out” over reward-seeking behavior in all conditions. If an animal is starving, it would be adaptive to expose itself to potential dangers in order to obtain food and survive. In this case, one would predict that food restricted animals may be less sensitive to external cues predicting danger. In fact, previous work has shown that calorically restricted animals show reduced fear and anxiety-related behaviors as a result of hindbrain (Maniscalco et al., 2015) and hypothalamic (Padilla et al., 2016) signals to forebrain nuclei including the amygdala. Hunger has also been explicitly demonstrated to suppress competing motivation systems including anxiety and fear (Burnett et al., 2016). For these reasons, we hypothesized that the interactions between BLA neurons would shift in hungry mice.

Following food deprivation, we found that BLA→NAc neurons were no longer inhibited by BLA→CeM photostimulation, but rather this manipulation lead to increased current evoked firing in these cells. Moreover, BLA→NAc neurons were also more sensitive to the activation of neighboring BLA→NAc neurons in slices taken from food deprived mice. These satiety-dependent changes in the interactions among valence coding BLA neurons could potentially facilitate the acquisition of caloric rewards by hungry animals despite the presence of risks.

Notably, although both populations we studied here modulated firing in the opposing population, neither *directly* evoked firing in the other during light-only stimulation. If this were not the case, activation in BLA→NAc neurons might recruit a negative feedback loop whereby their activity would be shunted by BLA→CeM neurons in satiety, and during food deprivation, the reciprocal excitatory relationship between these populations could lead to epileptogenesis. Instead, in both satiety and food deprivation, BLA→NAc neurons appear to propel BLA→CeM cells into a state primed for activation, in which additional direct stimulation of the latter results in robust BLA→CeM neurons firing. Such a relationship may be particularly advantageous in helping to prepare animals to detect threats during reward seeking, when they may be at relatively greater vulnerability to risk.

The finding that BLA→NAc neurons exerted no significant net effect upon the activity of neighboring BLA→ NAc neurons in satiety was somewhat surprising; we predicted that there would be a largely excitatory relationship between cells of the same projection-target identity, and in fact, we *did* observe within-population potentiation of firing among BLA→CeM neurons. It is likely that the diverse impacts of BLA→NAc activation upon other BLA→NAc neurons in slices from sated mice stems from the marked heterogeneity of valence coding functions within the NAc itself (Al-Hasani et al., 2015; Ambroggi et al., 2011; Castro and Berridge, 2014; Faure et al., 2010; Kravitz et al., 2012; Lobo et al., 2010; Namburi et al., 2016; Roitman et al., 2005). It is also possible that NAc projecting BLA neurons represent a population composed of multiple subgroups of functionally-defined cells. In fact, we observed heterogeneous response profiles in both BLA→ NAc and BLA→CeM neurons during presentation of sucrose- and quinine-predictive cues *in vivo* (Beyeler et al., 2016), underscoring the complexity of valence coding in these populations. That BLA→NAc neurons within-population stimulation *did* lead to significantly increased firing following food deprivation suggests that there is unity among this population given a considerable motivational drive, so it is possible that this population functions as a concordant group only under particular conditions.

### Understanding previous observations through the lens of local interactions in the BLA

Our current findings clarify some open questions arising from our previous work. We have reported that photoinhibition of BLA→CeM neurons during the pairing of the conditioned stimulus and footshock in the acquisition phase impaired fear learning (Namburi et al., 2015), highlighting the necessity of this population to negative valence encoding. Importantly, inhibition of BLA→CeM neurons during reward learning *enhanced* performance in this task (Namburi et al., 2015), consistent with the present finding that BLA→CeM cells inhibit reward-encoding BLA→NAc neurons. Photoinhibition of BLA→NAc neurons did not similarly modulate fear and reward learning (Namburi et al., 2015), a negative result which may be partially explained by the lack of inhibition of BLA→CeM neurons by BLA→NAc neurons in either the sated or the food deprived conditions. As BLA→NAc neurons fail to inhibit BLA→CeM neurons, it is unsurprising that inhibiting this population failed to potentiate fear learning; BLA→CeM neurons would not have been disinhibited by optical inhibition of BLA→ NAc neurons.

We have also previously reported that BLA→CeM neurons inhibit proportionally more neighboring BLA neurons than BLA→NAc neurons in intact animals (Beyeler et al., 2018). We collected these data from phototagging sessions at the end of a behavioral task in which the mice received large quantities of a 30% sucrose solution, to the point that they were sated. Given our *ex vivo* findings that BLA→CeM neurons unidirectionally inhibited firing in BLA→NAc cells in sated mice, it stands to reason that BLA→NAc neurons were among the cells inhibited by BLA→CeM neurons photostimulation *in vivo.* This ability of BLA→CeM neurons to powerfully inhibit neighboring cells - and thereby commandeer the activity of the BLA in sated animals - is congruent with the notion that fear learning can occur after a single pairing of a predictive cue with an aversive outcome (Fanselow, 1990). Contrastingly, we have reported that relatively few BLA neurons were inhibited by BLA→NAc photostimulation *in vivo,* consistent with the relative difficulty of reward learning, which requires multiple trials. As we observed diverse within-population interactions among BLA→NAc neurons in slices taken from sated mice, it is possible that some of the neurons we observed to be inhibited by BLA→ NAc photostimulation *in vivo* were neighboring, ChR2-negative BLA→NAc neurons. Alternatively, these photoinhibited cells could fall into a distinct subpopulation, such as projections to other downstream targets (McGarry and Carter, 2017), or local interneurons (Lucas et al., 2016). Together, our observations *in* and *ex vivo* suggest that BLA→CeM neurons uniquely inhibit neighboring BLA neurons, potentially to facilitate fast recruitment of escape behaviors in cases of immediate threat. However, it remains to be tested whether the capability of BLA→CeM neurons to suppress activity in BLA→NAc neurons extends to other populations of BLA neurons, and whether other populations of BLA neurons more closely resemble BLA→ NAc or BLA→CeM neurons in their local interactions.

The local interactions in the BLA we observed here also help to reconcile our previous observations to disparate results in the literature. Previous studies have reported that animals will form a real time place preference for the side of an arena paired with optical stimulation of BLA terminals within the NAc (Britt et al., 2012), will robustly self-stimulate this pathway (Stuber et al., 2011), and stimulation of this pathway can mitigate a depression-like state (Ramirez et al., 2015). However, in our hands, mice did not form a place preference and only modestly performed intracranial self-stimulation of BLA→ NAc neurons (Namburi et al., 2015). The subtle differences in our findings may be accounted for by the local interactions among BLA neurons populations described here, as well as the placement of the optic fiber and viral vectors used in these three studies. To stimulate BLA→NAc cells, we placed the optical fiber above BLA and delivered stimulation to somata, which could have promoted enhanced activity in BLA→CeM neurons and ultimately diluted any positive valence signaled by the BLA→NAc neurons. In the pioneering studies (Britt et al., 2012; Stuber et al., 2011), the optic fiber was implanted above the NAc. Such terminal stimulation may have circumvented local interactions between neuronal populations within the BLA, resulting in a more coherent, uncontested positive valence signal.

### Food deprivation primarily alters activity in BLA→NAc neurons

Using two-photon imaging of GCaMP6m activity, we observed that food deprivation impacted calcium transients in both neurons populations that we studied. However, we did not detect a notable impact of food deprivation upon BLA→CeM neurons *ex vivo,* either in the impact of BLA→NAc stimulation upon these neurons or the recorded amplitude and frequency of mEPSCs. It is likely that the discrepancy between these measures stems from the loss of broad-level connectivity that occurs in the slice preparation. However, we were able to measure consistent satiety-state dependent changes in BLA→NAc neuron activity in every approach that we used.

Considered together, our findings suggest that the increased frequency of calcium transients in BLA→NAc neurons that we observed in intact, food deprived mice arise from a change in afferent input to these cells. We did not observe robust changes in the intrinsic properties of BLA→NAc neurons following food deprivation, suggesting that the increased activity we measured in these cells was unlikely to have been caused by a change in membrane excitability. We did, however, observe an increase in the frequency of mEPSCs onto BLA→NAc neurons in slices from food deprived mice, which suggests a commensurate increase in presynaptic release probability. Although we cannot rule out an anatomical change (e.g., an upregulation of glutamatergic synapses onto BLA→NAc neurons during food deprivation, consistent with Wable et al., 2014), we suspect increased activity in glutamatergic afferents specifically targeting BLA neurons projecting to the NAc. In addition to an increase in excitatory presynaptic signaling, the increased calcium activity among BLA→NAc neurons we observed *in vivo* could potentially stem from a release in feed-forward inhibition from BLA→CeM neurons, as this population underwent a concurrent reduction in activity during food deprivation.

Because the state-dependent changes we observed primarily manifested as an increase in BLA→ NAc activity, we tested the hypothesis that this projection-specific alteration was necessary for invigorating reward-seeking behavior in hungry mice. We found that BLA→NAc inhibition did not impact food intake when standard chow was readily accessible, but when the animals were required to work for a palatable reward, this same inhibition reduced the appetitive motivation of food deprived mice. These findings suggest that positive valence encoding BLA neurons are not critical for feeding *per se,* but activity in this population might drive foraging and reward-seeking behaviors to increase the likelihood that animals obtain food rewards.

In addition to elucidating the sensitivity of valence processing neurons in the BLA to satiety-state, these data also imply a cautionary point regarding the prudence of food regulation as a means of increasing the willingness of experimental animals to perform behavioral tasks. Whereas one might expect food restriction to impact the neural systems responsible for the detection and maintenance of energy balance, the influence of satiety upon other neural systems often escapes consideration. Given the marked impact of food deprivation upon BLA projection neurons that we observed here, as well as the increasingly apparent interconnectivity of all brain systems, it is likely that food restricting animals for behavior influences the systems under study, even if they are ostensibly unrelated to feeding.

Here, we characterize the impact of changes in satiety upon the activity and local interactions of two populations of BLA projection neurons oppositely encoding valence, and provide a novel mechanism that may flexibly drive action selection depending upon the internal needs of the animal. Understanding how valence encoding systems interact at baseline and in food deprived conditions provides a foundation for measuring how other experiences impact microcircuit dynamics in the BLA. Moreover, establishing this fundamental understanding of the relationship between emotional valence processing and satiety enables the discovery of how these systems may become dysfunctional in psychiatric conditions in which both valence processing and energy homeostasis become disrupted, such as eating disorders and depression.

## Methods

### Animals and Stereotaxic Surgery

We used 93 adult wild-type C57BL/6J male mice (8-10 weeks old, Jackson Laboratory, Bar Harbor, ME) for the experiments presented in this manuscript. After surgery, animals were housed in reverse 12 hour light/dark cycles with free access to food and water for at least 10-12 weeks to allow for viral expression. Animals were handled under procedures in accordance with the guidelines from the NIH and with approval from the MIT Committee on Animal Care (CAC). All surgeries were conducted in aseptic conditions with a digital small animal stereotaxic instrument (David Kopf Instruments, Tujuna, CA).

Mice were anaesthetized with isoflurane (5% for induction, 1.5-2.0% afterward) in the stereotaxic frame for the duration of the surgery and their body temperatures were maintained with a heating pad. To restrict expression of ChR2-eYFP to basolateral amygdala (BLA) neurons projecting to a specific downstream target, we injected an adeno-associated virus serotype 5 (AAV5) carrying channelrhodopsin-2 (ChR2) fused to an enhanced yellow fluorescent protein (eYFP) in a double-floxed inverted open reading frame (DIO) under the control of elongation factor-1α promoter (∼400 nl of AAV5-EF1α-DIO-ChR2-eYFP) into the BLA at stereotaxic coordinates from bregma: −1.60 mm AP, +3.35 mm ML and −5.00 mm DV. Concurrently, canine adenovirus-2 (CAV2) carrying Cre-recombinase was injected into the nucleus accumbens (NAc, 300 nl), or the medial part of the central amygdala (CeM, 100 nl) using the following coordinates from bregma: NAc: 1.42 mm AP, +0.78 mm ML and −4.70 mm DV; CeM: −0.75 mm AP, +2.35 mm ML and −5.08 mm DV. In the animals used for *ex vivo* experiments, we also injected red retrogradely travelling fluorescent microspheres (RetroBeads, Lumafluor Inc.) into the NAc (60 nl) or CeM (40 nl). We used a similar protocol to achieve GCaMP6m expression in projection-target defined BLA neurons, in which we infused CAV2-Cre into the NAc or CeM and we injected 400 nl AAV5-CAG-flex-GCaMP6m into the right BLA. Finally, in the mice used for the progressive ratio behavioral experiment, we expressed inhibitory designer receptors exclusively activated by designer drugs (DREADDs) bilaterally in BLA→ NAc projection neurons by infusing 300 nl CAV2-Cre into the NAc of each hemisphere, as well as 400 nl AAV5-hSyn-DIO-hM4D(Gi)-mCherry bilaterally into the BLA. For mice in the control group, we instead infused 400 nl AAV5-hSyn-DIO-mCherry bilaterally into the BLA.

AAV5-EF1α-DIO-ChR2-eYFP virus was obtained from the University of North Carolina Vector Core (UNC, Chapel Hill, NC), and the CAV2 vector was obtained from the Kremer lab (CNRS, Montpellier, France). AAV5-hSyn-DIO-hM4D(Gi)-mCherry was purchased from Addgene and AAV5-hSyn-DIO-mCherry was acquired from UNC. The Genetically-Encoded Neuronal Indicator and Effector (GENIE) program at the Janelia Farm Research Campus (Loudoun County, Virginia) provided GCaMP6m, which was packaged by University of Pennsylvania Vector Core (UPENN, Philadelphia, PA).

All injections were performed using glass micropipettes (1-5 ml; Drummond Scientific) pulled with a puller (Narishige PC-10) and mounted in 10 ml microsyringes (Hamilton Microlitre 701; Hamilton Co.). Retrobeads and virus were delivered at a rate of 1 nl/s using a microsyringe pump (UMP3; WPI, Sarasota, FL) and controller (Micro4; WPI, Sarasota, FL). In all cases, the needle was raised 100 μm after completion of the injection and left for 10 minutes to allow the virus or retrobeads to diffuse from the injection site, and then slowly withdrawn. After retracting the injection needle in mice used for two-photon calcium imaging experiments, we implanted a 7.3mm gradient indexed (GRIN) microendoscope lens (0.5 NA, 3/2 pitch) with 0.6 mm diameter (Inscopix, Palo Alto, CA) above the right BLA at coordinates from bregma: −1.60 mm AP, +3.35 mm ML and −4.80 mm DV. We then placed a small aluminum head-bar (2cm x 2mm x 2mm) on the skull 1.5 mm anterior to bregma and cemented the lens and head-bar using a layer of adhesive cement (C&B metabond; Parkell, Edgewood, NY) followed by a layer of cranioplastic cement (Dental cement; Stoelting, Wood Dale, IL). After the cement dried, the GRIN lens was covered with a silicone gel (Kwik-Sil Adhesive, WPI, Sarasota, FL) to protect it between imaging sessions.

After surgery, each mouse’s body temperature was maintained with an infra-red heat lamp until he fully recovered from anesthesia. Slice electrophysiological recordings were conducted at least 10 weeks following surgery, and two-photon imaging began 12 weeks subsequent to surgery. Similarly, behavioral training started approximately 12 weeks after viral surgery.

### Two-photon calcium imaging of BLA→NAc and BLA→ CeM neurons

#### Measurement of Point-Spread Function

Similar to what is reported in Barretto et al., 2009, we used a 920 nm laser to image a 1 μm diameter fluorescent bead (TetraSpeck Fluorescent Microshpere Size Kit T14792; ThermoFisher, Waltham, MA) under a GRIN lens (0.5 NA) below the microscope objective (0.48 NA). We normalized the fluorescence signals to their maximum value in the cross-sectional profiles in the lateral and axial dimensions of the bead with varying distances. We fitted the resulting normalized fluorescence signals to a Gaussian distribution, and report the full-width at half maximum.

#### Pre-exposure of mice to palatable rewards

Mice were pre-exposed to a palatable, calorically dense reward (50% Ensure solution, diluted in water) in their home cage as well as in the experimental rig while being head-fixed. While head-fixed in the experimental rig, animals received 50% Ensure delivered randomly (drawn from an exponential distribution with an average ITI of 15 seconds) at a rate of 3.6μl/delivery. Prior to beginning testing, mice were acclimated to head-fixing over the course of a week, during which time imaging duration gradually increased from 15 minutes to 40 min.

#### Two-photon Microscopy

Each experimental imaging session lasted approximately 30-40 min. We first positioned the un-anesthetized animal in the head-fixed rig such that the GRIN lens was aligned with the air-immersed objective (20X, 0.48NA; Olympus, Center Valley, PA) of an upright two-photon (2P) fluorescence microscope (Ultima Investigator DL Laser Scanning Microscope; Bruker, Billerica, MA). We used wide-field epifluorescence imaging to visualize the tissue through the GRIN microendoscope and determined if GCaMP6m expression was sufficiently bright. After confirming the expression level, we used a Mai Tai DeepSee Ti:Sapphire 2P laser (Spectra Physics, Santa Clara, CA) at 920 nm wavelength emitting 64 mW at the level of the objective to image in Resonant Galvo scanning mode and collect GCaMP6m emission signals filtered through 525/70 nm with a photomultiplier tube (PMT). We acquired images of 512 x 512 pixels at an initial frame rate of approximately 30 Hz, then averaged every 4 images to yield a frame rate of approximately 7.5 Hz.

All mice were fed *ad libitum* prior to imaging in the sated condition during the dark portion of their light cycle on day 1 (session 1). We then food deprived the mice for 20 hours before imaging on day 2 (session 2). Afterwards, the animals were returned to *ad libitum* feeding, allowed two full days to return to the sated condition, and imaged again on day 4 (session 3). In each animal, we imaged multiple planes at least 50 μm apart (each for approximately 10 minutes), making note of the position of the objective relative to the GRIN lens for each plane in order to facilitate finding it in subsequent sessions. Individual planes were imaged in the same order on all three imaging sessions to ensure that any differences in Ca^2+^ activity observed across the sessions was not due to differing durations of head-fixing.

#### Analysis

GCaMP6m signals were collected using Prairie View software (Bruker, Billerica, MA) and then exported for analysis in MATLAB (MathWorks, Natick, MA). To obtain time traces of calcium activity, we first applied a non-rigid motion-correction algorithm (Suite2P; MATLAB) plus a rigid motion-correction algorithm (moco; ImageJ), and then calculated the relative fluorescence changes (ΔF/F) for the ROI of each cell. ROIs were manually fitted to the somata of GCaMP6m-expressing cells identified from the average image of the entire stack, as reported elsewhere (Bocarsly et al., 2015; McHenry et al., 2017; Otis et al., 2017). We excluded all ROIs for which there were no calcium transients across all three imaging sessions from our analysis (46 ROIs were excluded in total, leaving 95 active ROIs for analysis). The baseline fluorescence was defined as the 20th percentile of raw data. We manually counted the number of transient events within a session as a proxy for cellular activity.

### Ex vivo electrophysiology

#### Brain tissue preparation

Sated or food deprived mice were weighed and then anesthetized with 90 mg/kg pentobarbital and perfused transcardially with 10 mL of cold artificial cerebrospinal fluid (ACSF, at ∼4°C) containing (in mM): 75 sucrose, 87 NaCl, 2.5 KCl, 1.3 NaH2PO4, 7 MgCl2, 0.5 CaCl2, 25 NaHCO3 and 5 ascorbic acid. The brain was then extracted and glued (Roti coll 1; Carh Roth GmbH, Karlsruhe, Germany) to the platform of a semiautomatic vibrating blade microtome (VT1200; Leica, Buffalo Grove, IL). The platform was placed in the slicing chamber containing modified ACSF at 4°C. 300 μm coronal sections containing the NAc, CeM, and BLA were collected in a holding chamber filled with ACSF saturated with 95% O2 and 5% CO2, containing (in mM): 126 NaCl, 2.5 KCl, 1.25 NaH2PO4, 1.0 MgCl2, 2.4 CaCl2, 26.0 NaHCO3, and 10 glucose. Recordings were started 1 h after slicing, and the temperature was maintained at approximately 31°C both in the holding chamber and during the recordings.

All viral injection sites were checked and imaged with a camera (Hamamatsu Photonics K.K., Japan) attached to the microscope (BX51; Olympus, Center Valley, PA). The slice images were registered to the mouse brain atlas (Paxinos and Watson) and the brightest point of fluorescence was considered the center of the injection. Data were not collected from animals with injection sites outside the BLA, NAc, or CeM.

#### Whole-cell patch-clamp recording

Recordings were made from visually identified neurons containing red retrobeads, with the exception of those recordings used to verify DREADDs functionality, which were made from neurons visually confirmed to express mCherry. Recorded neurons were filled with Alexa Fluor (AF) 350 and biocytin. Voltage and current-clamp recordings of BLA projection neurons were conducted using glass microelectrodes (3-8 MΩ) molded with a horizontal puller (P-1000, Sutter, CA). Recorded signals were amplified using a Multiclamp 700B amplifier (Molecular Devices, Sunnyvale, CA). Analog signals were digitized at 10 kHz using a Digidata 1440 and recorded using the pClamp10 software (Molecular Devices, Sunnyvale, CA). Oxygenated ACSF was perfused onto the slice with a peristaltic pump (Minipuls3; Gilson, Middleton, WI) at ∼3 mL/min. Cells that lacked a constant inward current in response to a 1 s constant blue light pulse in voltage-clamp were confirmed to lack ChR2 expression.

#### Voltage-clamp experiments

The microelectrodes were filled with a solution containing (in mM): 120 caesium methansulphonate, 20 HEPES, 0.4 EGTA, 2.8 NaCl, 5 tetraethylammonium chloride, 2.5 MgATP, 0.25 NaGTP, 8 biocytin and 2 Alexa Fluor 350 (pH 7.3, 287 mOsmol). Cells were held at −70 mV holding potential to record excitatory currents and at 0 mV holding potential to record inhibitory currents in response to 5 ms, 5 mW blue light pulses delivered every 10 seconds via a 470 nm LED light source (pE-100; CoolLED, NY, USA). In order to determine whether responses were monosynaptic, the optically-evoked currents were recorded in the presence of tetrodotoxin (TTX; 1μM) and 4-Aminopyridine (4AP; 100 μM). Additionally, in a subset of cells, we bath applied 100 mM of the gamma-aminobutyric acid receptor A (GABAaR) antagonist picrotoxin (PTX; R&D systems) during recording of the current at 0 mV to determine whether it was mediated by GABAaR. Finally, to determine whether the current at −70 mV was mediated by α-amino-3-hydroxy-5-methyl-4-isoxazolepropionic acid receptors (AMPAR), we bath applied 20 μM 2,3-dihydroxy-6-nitro-7-sulfamoyl-benzo[f]quinoxaline-2,3-dione (NBQX; Tocris Bioscience, Bristol, UK). In a separate experiment, we held cells at −70 mV holding potential in the constant presence of 1μM bath TTX to enable continuous recording of miniature excitatory post synaptic currents (mEPSCs).

#### Current-clamp experiments

The microelectrodes were filled with a solution containing (in mM): 125 potassium gluconate, 20 HEPES, 10 NaCl, 3 MgATP, 8 biocytin and 2 Alexa Fluor 350 (pH 7.33; 287 milliOsmol). The amount of somatic current required to drive stable firing near 3-5 Hz was calculated for each cell using progressively increasing current steps. <u>Stimulation protocol:</u> Firing was evoked using a square pulse of somatic current injection between 50 and 300 pA for 7.5 s every 30 s. Light ON and light OFF sweeps were interleaved, such that every other sweep included delivery of a 7.5 s, 10 Hz train of 5 ms light pulses with a light power density of 14.2 mW/mm^2^ (4.4 mW with 40x objective) delivered via a 470 nm LED light source. The timing of the LED stimulation and current injection were offset so that the LED train began 2.5 s before current injection. For the verification that Clozapine N-oxide (CNO) action at the hM4Di receptor inhibits cells, we recorded the responses of hM4Di- or mCherry-expressing cells to a 300 pA current ramp and to square current steps of increasing amplitude (starting at 50 pA with a Δ of 25 pA), as well as continuous activity before and after bath application of 10 μM CNO (Tocris Bioscience, Bristol, UK). These experiments were performed in 100 mM PTX.

#### Data analysis

Off-line analysis was performed using Clampfit software (Molecular Devices, Sunnyvale, CA), as well as MiniAnalysis Version 6 (Synaptosoft, Decatur, GA) and MATLAB (MathWorks, Natick, MA) using Matlab codes written by A.M.L. and P.N. For voltage-clamp experiments, the average amplitudes and onset latencies of excitatory and inhibitory currents were measured from the onset of the light pulse. mEPSCs were measured across the first three minutes of holding current stability, and were included in the analysis based upon meeting the following five criteria: (1) the decay was longer than the rise time, (2) the amplitude was at least two times the standard deviation of the baseline, (3) the rise time was less than 7 ms, (4) the decay could be fitted by a decaying exponential function, and (5) the ratio of the amplitude to the rise time was greater than 1.

For current-clamp experiments, average firing rate in light ON and light OFF conditions was measured during the first 5 s of the current pulse, corresponding to the epoch during which light stimulation and current coincided. Additionally, the firing rate during the first 2.5 s of LED stimulation in the light first protocol was measured as the “light only” response, and the firing rate in the 2.5 s preceding LED stimulation was measured as “basal” activity. Resistance was measured in voltage clamp as the difference between holding current required to clamp the cell membrane to −70 mV and the mean current at the last 5 ms of a 5 mV voltage pulse lasting 100 ms. For each cell, a minimum of 10 pulses were averaged. Spike properties were calculated based on the first spike elicited by the slow ramp protocol performed in current clamp. Spike threshold was measured as the first voltage point in which the derivative was greater than 1.5 mV/sec. Spike amplitude was calculated as the maximum voltage during the spike minus the spike threshold voltage. Spike halfwidth was measured as the time difference between the half-amplitude point in the up-stroke and down-stroke of the action potential. The rheobase was calculated using the slow ramp protocol, and was measured as the current at the first spike threshold. The relationship between current steps and spike frequency was recorded in current clamp. The responses of each cell were fitted to the following expression: f = A1 - A2*exp(-I/tau_i), and maximal frequency was measured as the maximal number of spikes recorded for a given cell. For verification of DREADDs functionality, we measured average membrane potential values across two minutes leading up to the application of 10 μM CNO, and compared those values to average membrane potential values across two minutes beginning two minutes after the application of CNO.

### Progressive Ratio and Feeding Behaviors

#### Feeding task

We used a within-subjects design in which every mouse received intraperitoneal (i.p.) injections of 1 mg/kg clozapine n-oxide (CNO, Tocris Bioscience, Bristol, UK) or vehicle (0.9% sterile saline) separated by one week and counterbalanced for order. The feeding task was conducted at the beginning of the dark phase of the light/dark cycle, starting at 9 AM. In preparation for the experiment, mice were singly housed and food deprived at lights on (9 PM) the previous night. CNO or vehicle was administered at 8:30 AM, half an hour before one pellet of standard chow was weighed and made available to each mouse in his home cage at 9 AM. We then weighed the pellet at 10 AM (+1 hr), 11 AM (+2 hrs), and finally at 1 PM (+4 hrs) to determine how much chow each mouse consumed. After the +4 hrs measurement, we returned the mice to grouped housing with their previous cage mates, and provided food *ad libitum*.

#### Progressive ratio training

All progressive ratio training and testing was conducted in operant chambers (MedAssociates, Fairfax, VT) placed in custom-made sound-attenuating boxes. Each chamber contained one nose-poke port with an infrared beam break, and one liquid reward delivery port with a house-light positioned above it, all located on the same side of the chamber. Delivery of a 15μl bolus of a palatable, calorically dense reward (Ensure, diluted to 50% in water) into the reward port coincided with illumination of the house-light (extinguished when the mouse retrieved the reward or after 10 sec, whichever occurred first), and was contingent upon the mouse performing the requisite number of nose pokes.

Mice were allowed *ad libitum* access to food for the duration of operant training, and were not food deprived until testing. They were first trained on a fixed ratio 1 (FR1) schedule, in which each nose poke into the operant port resulted in the delivery of one bolus of 50% Ensure. Training on the FR1 schedule lasted at least five days, or until the mice made at least 200 nose pokes in the two hour session (no more than 8 days). Subsequently, we trained the mice on a progressive ratio (PR) schedule, in which the delivery of each reward required more effort than the previous one. The PR schedule was determined using the following equation, as described in Richardson and Roberts, 1996:

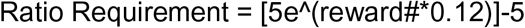

This equation resulted in the following series of ratio requirements (listed here up to 40 reward deliveries): 1, 1, 2, 3, 4, 5, 7, 8, 10, 12, 14, 16, 19, 22, 25, 29, 33, 38, 44, 50, 57, 65, 74, 84, 95, 108, 123, 139, 157, 178, 201, 228, 257, 291, 328, 371,419, 473, 534, 603, etc. Training on the PR schedule lasted at least five days, or until mice achieved at least 24 reward deliveries (requiring 623 total nose pokes) in the three hour session.

#### Progressive ratio testing

All animals were food deprived for 20 hours prior to testing, which was always conducted in the food deprived state. 30 minutes prior to beginning the PR session, animals were administered either 1 mg/kg CNO (in a 0.2 mg/ml solution) or vehicle (0.9% saline solution) intraperitoneally.

We used a within-subjects design, and each mouse was tested twice in the CNO condition and twice in the vehicle condition (four total test days). We counterbalanced the order in which animals received CNO and vehicle, always allowing at least two days to pass between test sessions to be sure that drugs would be fully metabolized and washed out between test days. Mice were tested on the same PR schedule on which they were trained, and were refed *ad libitum* at the completion of the each test session.

#### Analysis

Behavioral responses (nose pokes and entries into the reward delivery port) and task events (reward deliveries) were recorded using MED-PC software (MedAssociates), and were analyzed offline using MATLAB codes written by P.N. and A.M.L. We used a 30 minute cut off requirement between reward deliveries, such that if more than 30 minutes passed between reward deliveries, we considered the session to have ended at the reward delivery prior to the pause in responding.

### Histology

#### Histology following two-photon imaging or progressive ratio behavior

After the completion of imaging or progressive ratio testing, the mice were anesthetized with pentobarbital sodium and transcardially perfused with ice-cold Ringer’s solution and then ice-cold 4% paraformaldehyde (PFA) in PBS (pH 7.3). Extracted brains were fixed in 4% PFA overnight then equilibrated in 30% sucrose in PBS solution. 40 μm thick coronal sections were sliced using a sliding microtome (HM430; Thermo Fisher Scientific, Waltham, MA) and stored in PBS at 4°C until they were processed for histology. Sections were then incubated with a DNA-specific fluorescent probe (DAPI: 4’,6-Diamidino-2-Phenylindole (1:50,000)) for 30 minutes and afterwards washed with PBS-1X then mounted on microscope slides with polyvinyl alcohol mounting medium with DABCO (Sigma, MO, USA).

#### Histology following slice electrophysiological recording

The location of all recorded neurons was confirmed after the recording. Co-localization of AF 350 and red retrobeads was confirmed with confocal microscopy for the cells that were recovered with biocytin-streptavidin staining. For each experiment, the slices containing a retrograde viral or retrobead injection (NAc or CeM) or a recorded neuron (BLA) were fixed overnight at 4°C in 4 % PFA, and then kept in PBS. Slices containing patched neurons were incubated for 2 hours in streptavidin-CF405 (2 mg/ml, dilution 1:500, Biotium, Hayward, CA), mounted on microscope slides with PVA-DABCO and imaged under the confocal microscope. All other slices were stained for a DNA-specific fluorescent probe (DAPI; 1:50,000). Following staining, all slices were mounted onto glass slides and coverslipped using PVA-DABCO (Sigma, MO, USA).

### Microscopy

Confocal fluorescence images were taken with an Olympus FV1000 confocal laser scanning microscope using a 20X, 0.75NA objective for the injection sites and 40X, 1.3NA objective for the recorded neurons. Viral injection and recording electrode placement images were acquired using the FluoView software (Olympus, Center Valley, PA). The center of the viral injection was chosen as the brightest fluorescent point in AP, ML and DV axes.

Histological verification of injection sites for mice used in the 2P Ca^2+^ imaging experiment and those used in the progressive ratio and feeding behaviors was performed using a BZX Fluorescence Microscope (Keyence Corporation of America, Elmwood Park, NJ) using a CFI Plan Fluor DL 10X objective (Nikon, Tokyo, Japan). Injection and GRIN lens placement images were acquired using BZX Analyzer software with the BZX Multi-Stack Module (Keyence Corporation of America, Elmwood Park, NJ). As with confocal images, the center of the viral injection was chosen as the brightest fluorescent point in AP, ML and DV axes.

### Statistical analysis

Statistical analyses were performed using MATLAB or a commercial software (GraphPad Prism; GraphPad Software, Inc, La Jolla, CA), and values are reported as mean ± standard error of the mean (S.E.M.). Data in all groups were evaluated for normality using a Kolmogorov-Smirnov test with Dallal-Wilkinson-Lilliefors p value. Data obeying a Gaussian distribution were compared using two-tailed paired Student t-tests (non-directional), or with one- or two-way ANOVA when three or more groups were compared. Nonparametric data were compared using Wilcoxon signed-rank tests, Mann Whitney U tests, or Rank Sum tests. Threshold for significance was placed at ^+^*p*<0.1, \**p*<0.05, \*\**p*<0.01 and \*\*\**p*<0.001).

---

**Figure S1.**
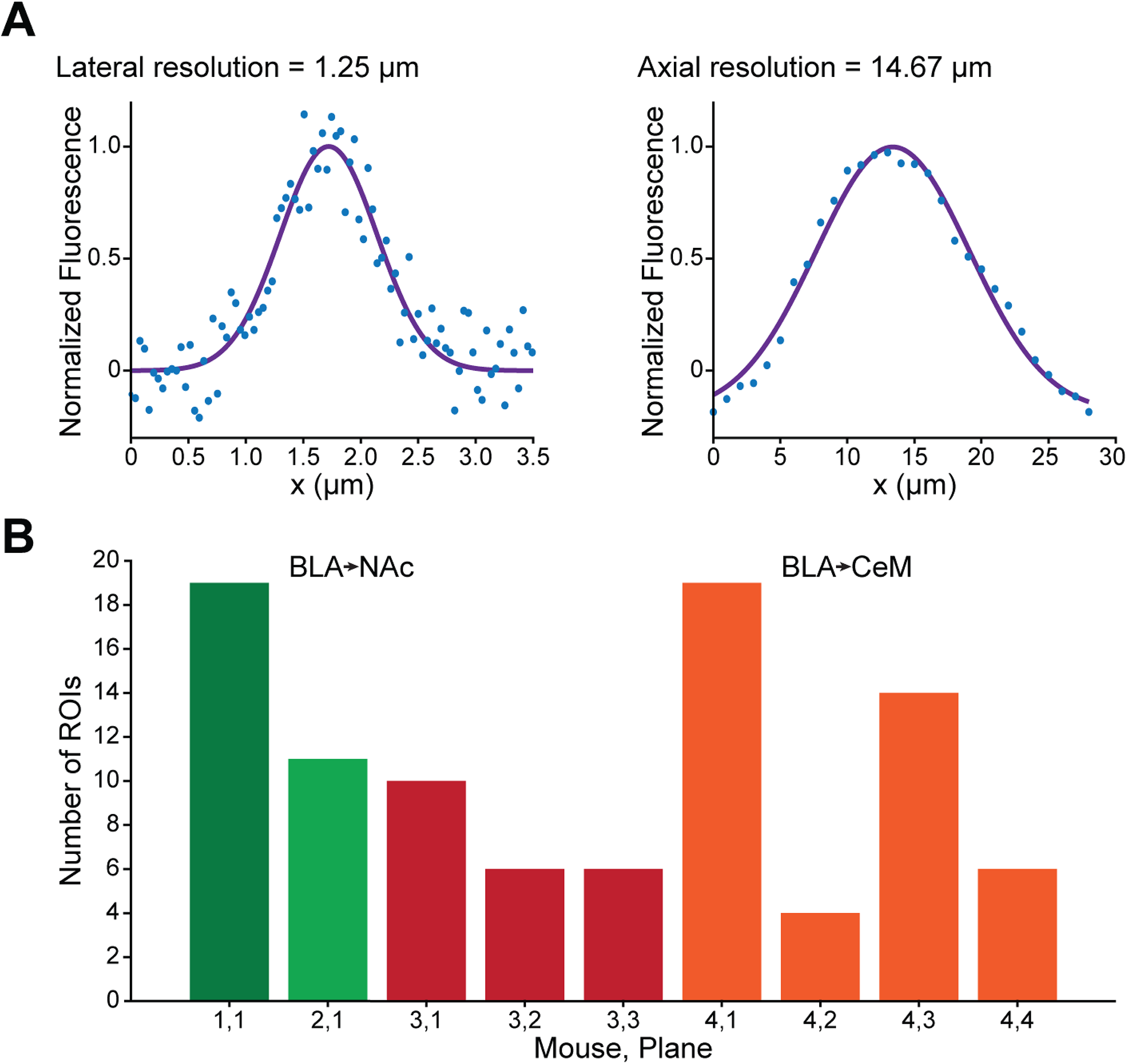
Calibration of two-photon signals (Relates to Main Figure 1) (A) Point-spread function in the lateral (left panel) and axial (right panel) planes. To evaluate the resolution of our two-photon measurements, we sampled the fluorescence signals in the cross-sectional profiles of a 1 μm diameter fluorescent bead under a GRIN lens below the microscope objective. The normalized signals (blue circles) were fitted with a Gaussian curve (purple line), and the full-width at half maximum of the lateral and axial profiles were 1.25 μm and 14.67 μm, respectively. (B) Number of ROIs per plane in the mice imaged across satiety and food deprivation. Each mouse is represented in a different color (BLA→NAc mice in greens, BLA→CeM mice in red and orange), and each plane in an individual bar.

**Figure S2.**
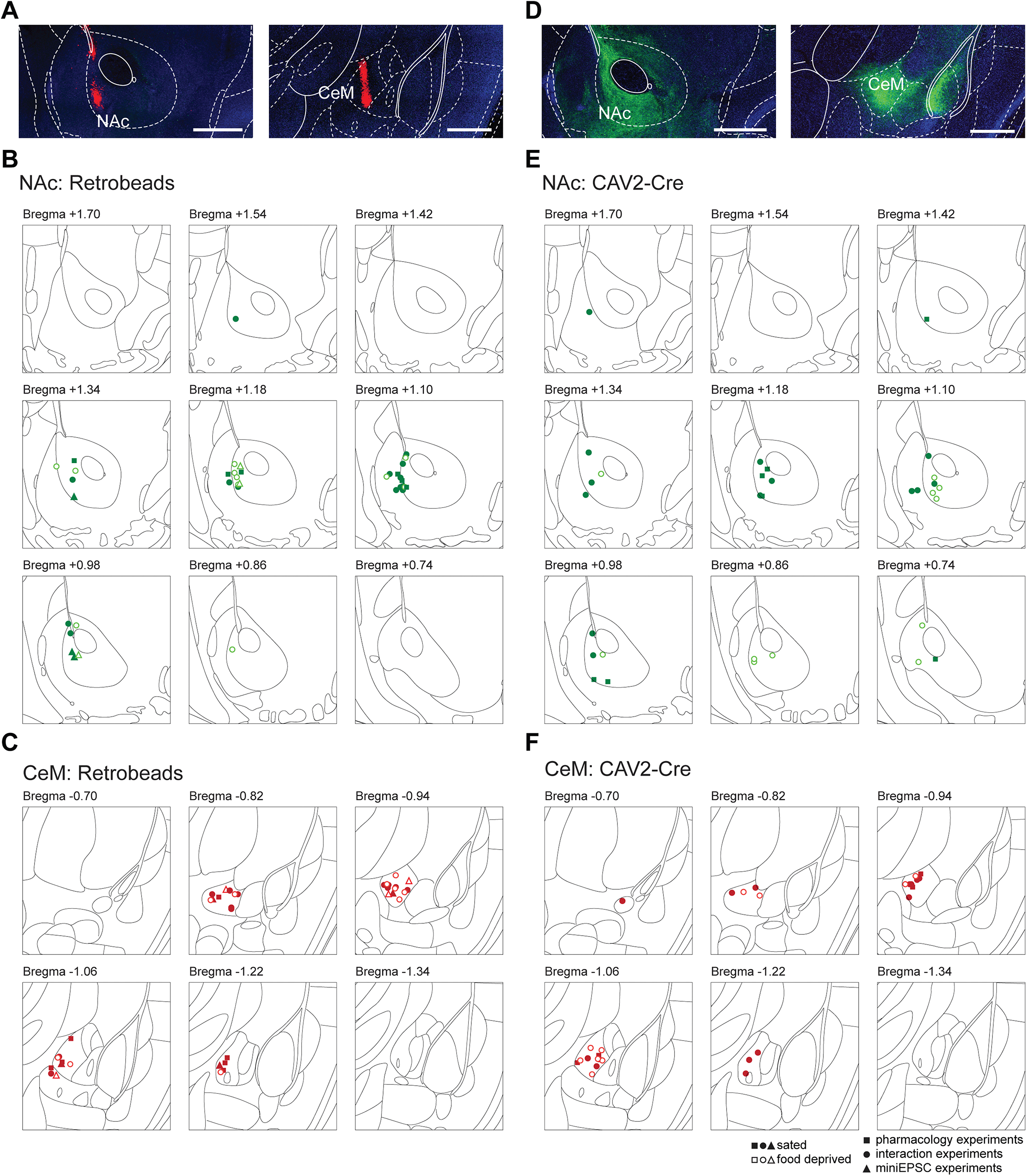
Injection sites for mice used in whole-cell patch-clamp experiments (Relates to Main Figures 2 and 3, and Supplemental Figures 3-7) (A) Representative confocal images showing red retrobead injection sites in the NAc (left) and CeM (right). Red = retrobeads, blue = DAPI, scale bars = 500 μm. (B) Center point of retrobead injection sites in the NAc. (C) Center point of retrobead injection sites in the CeM. (D) Representative confocal images showing ChR2-positive terminals of BLA neurons projecting to the NAc (right) and CeM (left). Green = ChR2-eYFP, blue = DAPI, scale bars = 500 μm. (E) Center point of CAV2-Cre injection sites in the NAc. (F) Center point of CAV2-Cre injection sites in the CeM. Filled markers = sated animals; open markers = food deprived animals. Squares = animals used in voltage clamp pharmacology experiments (supplemental figure 3), circles = animals used in current clamp microcircuit interaction experiments (main figure 2 and supplemental figures 4 and 5), triangles = animals used in mEPSC experiments (main figure 3).

**Figure S3.**
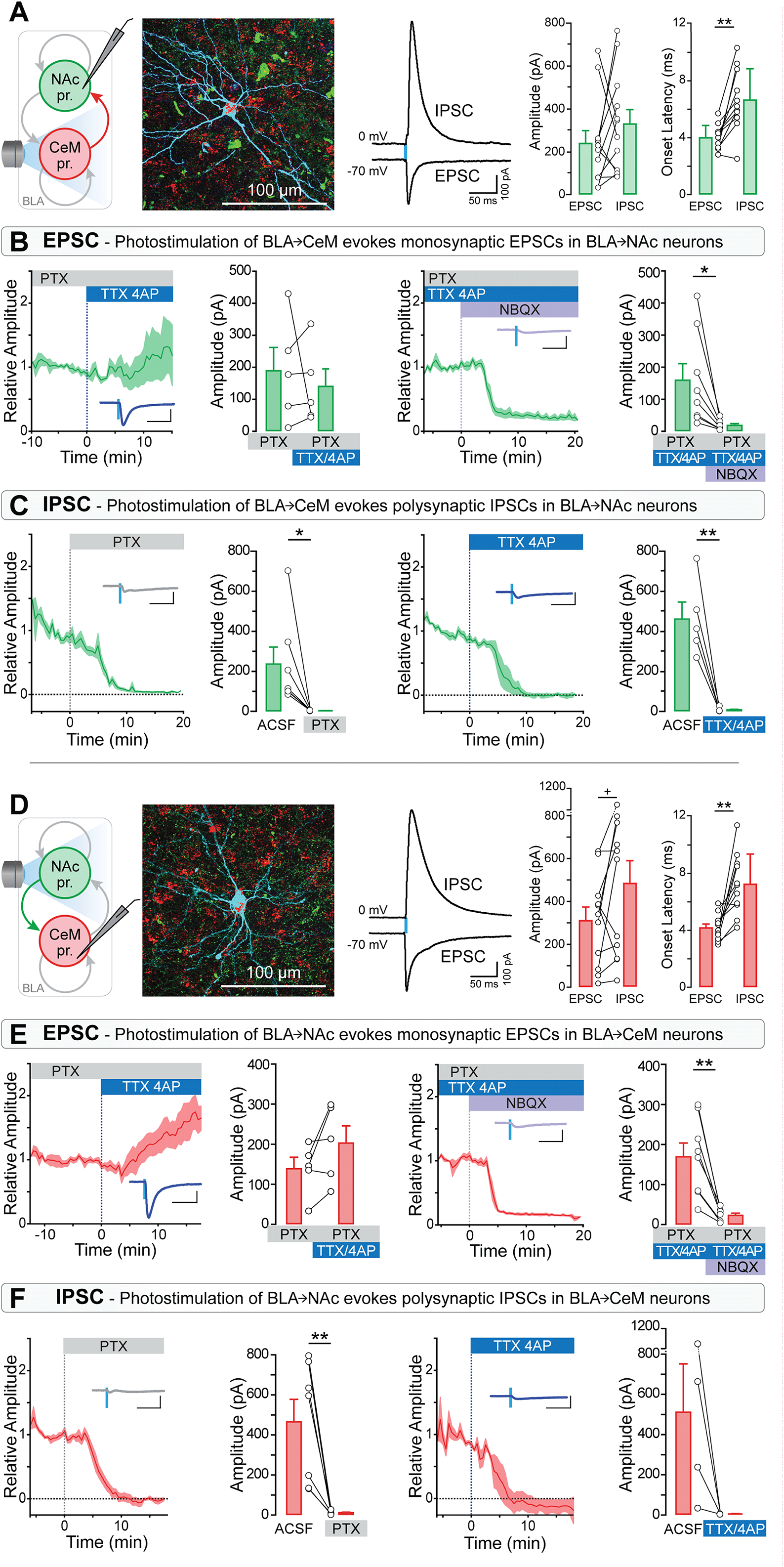
Microcircuit connectivity of BLA→NAc and BLA→CeM neurons (Relates to Main Figure 2) (A) We used red retrobeads to visually identify and patch NAc-projecting BLA neurons in order to measure synaptic responses to photostimulation of BLA→CeM neurons, and recorded 12 cells from 6 mice. Left panels: schematic diagram of experimental approach, and confocal image of a retrobead-positive recorded cell, processed for biocytin immunoreactivity (green = ChR2-eYFP, red = retrobeads, cyan = biocytin, scale bar = 100 μm). Middle panel: average EPSC and IPSC responses from a representative neuron, recorded at −70 mV and 0 mV, respectively; photostimulation of BLA→CeM neurons evoked EPSCs and IPSCs in all recorded BLA→NAc neurons. Right panels: The amplitudes of EPSCs and IPSCs in all recorded cells did not significantly differ (Wilcoxon signed-rank test, *p*=0.3804), but the onset latency of the EPSC was shorter than that of the IPSC (paired t-test, t_10_=4.540, \*\**p*=0.0011). (B) Photostimulation of BLA→CeM neurons evoked a monosynaptic AMPAergic EPSC in BLA→NAc neurons. The EPSC was blocked by neither PTX nor TTX/4AP (time course of normalized responses and average amplitude, left two panels; an example EPSC is shown as an inset in the left most panel), but was inhibited by NBQX (right two panels) (paired t-test, t_7_=2.989, \**p*=0.0202). (C) The IPSC evoked by BLA→CeM neurons photostimulation was eliminated by PTX (paired t-test, t_6_=2.729, \**p*=0.0342) and TTX/4AP (paired t-test, t_4_=5.330, \*\**p*=0.0060). (D) In a separate set of experiments, we patched 11 retrobead-positive BLA→CeM neurons from 7 mice, and optically stimulated ChR2-eYFP expressing BLA→NAc neurons. Left panel: schematic of experimental strategy, and example of a biocytin-filled, retrobead-positive BLA→CeM neurons (green = ChR2-eYFP, red = retrobeads, cyan = biocytin, scale bar = 100 μm). Middle panel: representative average responses of a BLA→CeM neuron to BLA→NAc stimulation recorded at −70 mV and 0 mV. BLA→NAc photostimulation evoked EPSCs and IPSCs in all recorded BLA→CeM neurons. Right panels: We observed a trend towards IPSCs having a greater amplitude than EPSCs (Wilcoxon signed-rank test, ^+^*p*=0.0674). The onset latency of the EPSC was shorter than that of the IPSC (paired t-test, t_11_=4.289, \*\**p*=0.0016). (E) The EPSC was blocked by neither PTX nor TTX/4AP (left two panels), but was inhibited by NBQX (right two panels; paired t-test, t_7_=4.814, \*\**p*=0.0019). (F) The IPSC was sensitive to PTX (paired t-test, t_6_=4.078, \*\**p*=0.0065) and TTX/4AP (paired t-test, t_3_=2.123, *p*=0.1239).

**Figure S4.**
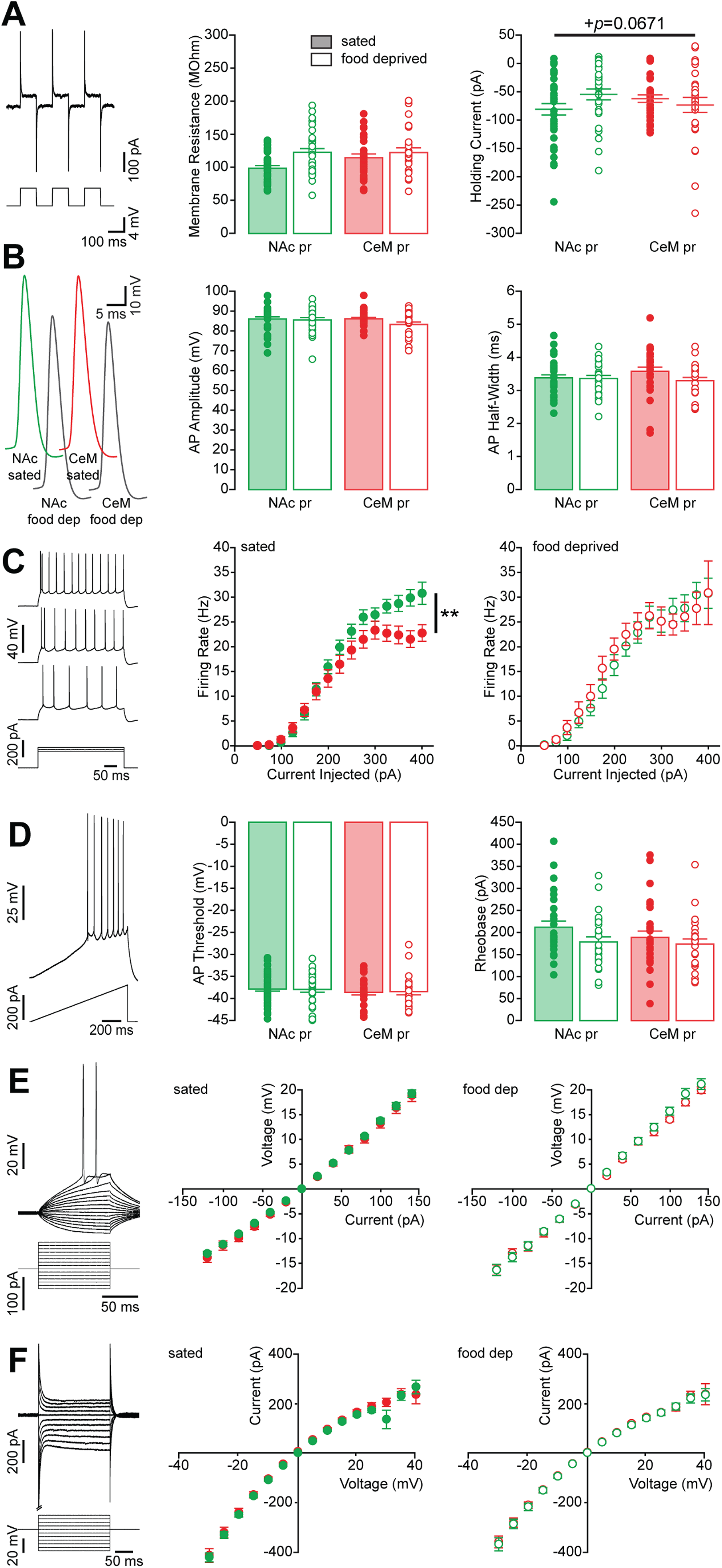
Membrane properties of BLA→NAc and BLA→CeM neurons (Relates to Main Figure 2) (A) Membrane resistance and holding current were calculated in all recorded cells based upon the current response to a 4 mV square pulse (example trace at left). Neither of these measures differed significantly between BLA→CeM and BLA→NAc neurons, nor were they impacted by satiety state (membrane resistance: two way ANOVA, projection*state interaction F_1,117_=1.861, *p*=0.1752; holding current: two way ANOVA, projection*state interaction F_1,117_=3.413, ^+^*p*=0.0671). (B) Example action potential traces recorded from BLA→NAc and BLA→CeM neurons from sated or food deprived mice (left panel). Action potential amplitude was not affected by the projection target nor by the satiety state (two way ANOVA, projection*state interaction F_1,112_=1.30, *p*=0.2567; center panel). Similarly, action potential half-width did not differ among the four conditions (two way ANOVA, projection*state interaction F_1,112_=1.4844, *p*=0.2256; right panel). (C) Voltage response of a cell to square pulse current injections of increasing magnitudes from a representative neuron (left panel). In slices taken from sated animals, BLA→CeM and BLA→NAc neurons responded to current injections with different maximum firing rates *(Mann-Whitney U=240.5,* \*\**p*=0.0019), however the current constant in the exponential fit of these curves (approximately 2/3 maximum firing rate) did not differ between BLA→NAc and BLA→CeM neurons *(Mann-Whitney U=342.0, p*=0.1257; center panel). The difference in maximum firing rate between the two projection target defined populations was abrogated following food deprivation *(Mann-Whitney U*=262.0, *p*=0.3501), and the current constant did not differ between BLA→NAc and BLA→CeM neurons in slices taken from food deprived mice *(Mann-Whitney U*=134.0, *p*=0.1572; right panel). (D) Representative example of a ramp current injection protocol used to determine the action potential threshold and rheobase, and corresponding voltage response (left panel). Action potential threshold did not differ among the four groups (center panel, two way ANOVA, projection*state interaction F_1,112_=0.0262, *p*=0.8717), nor did rheobase (center panel, two way ANOVA, projection*state interaction F_1,104_=0.4891, *p*=0.4859). (E) Representative example of the relationship between voltage and current recorded in current clamp mode (left panel); sweeps with action potentials were not considered in our analysis. There was no difference in the slope of the voltage/current relationship between BLA→NAc and BLA→CeM neurons from sated mice (center panel, unpaired t-test, t_42_=0.6899, *p*=0.4940), nor from food deprived mice (right panel, unpaired t-test, t_59_=0.2283, *p*=0.8202). (F) We also measured the relationship between current and voltage in voltage clamp mode, for which an example trace is presented at left. The resulting current/voltage curves are illustrated for BLA→NAc and BLA→CeM neurons in sated mice (center panel) and in food deprived mice (right panel).

**Figure S5.**
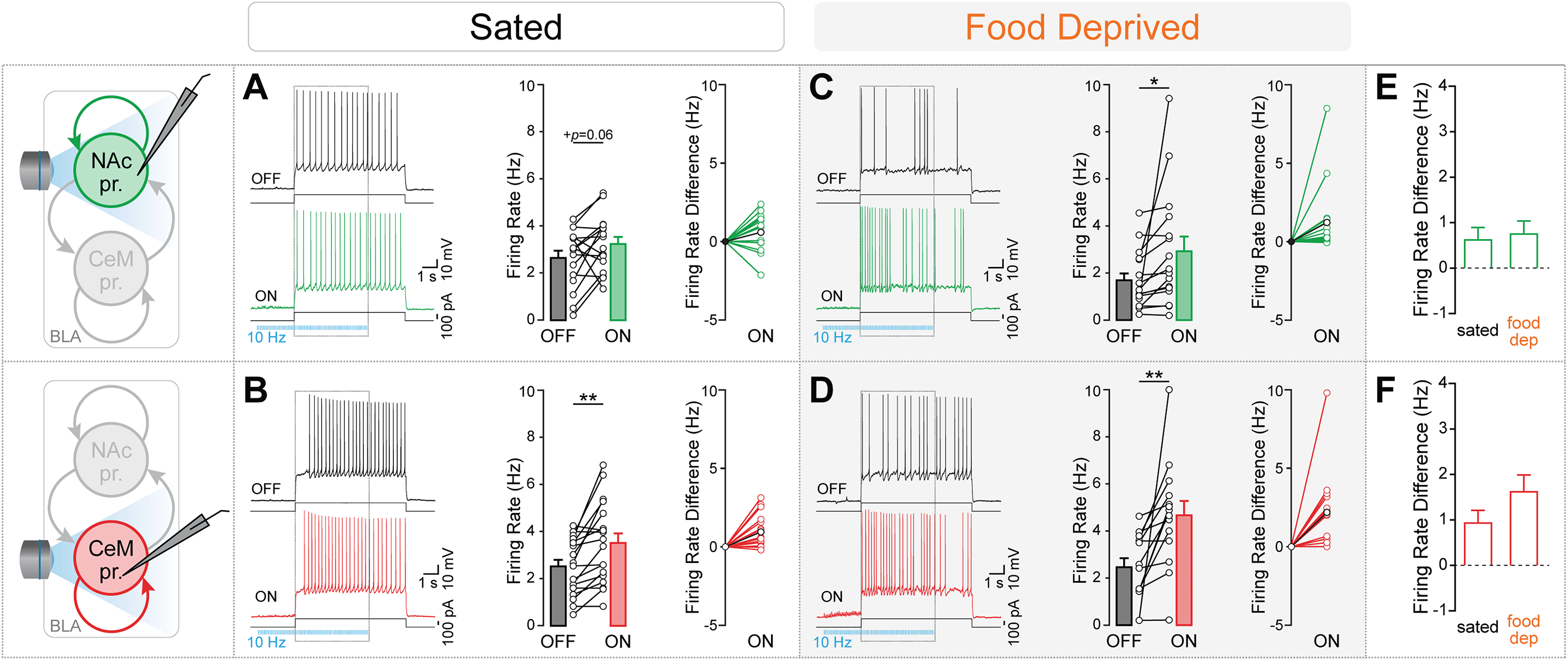
BLA within-population interactions do not dependent on satiety state (Relates to Main Figure 2) (A) Within-population photostimulation of BLA→NAc neurons did not evoke a inhibition of current-evoked firing in neighboring BLA→NAc neurons that did not express ChR2. Rather, retrobead-positive, ChR2-negative BLA→NAc neurons responded to 10 Hz photostimulation of neighboring ChR2-positive BLA→NAc neurons with heterogeneous changes in current evoked firing rate, trending toward an increase in firing rate (paired t-test, t_15_=2.033, ^+^*p*=0.0601). n=6 mice, 16 cells. (B) Within-population optical stimulation of BLA→CeM neurons in slices taken from sated mice increases current-evoked firing rate in neighboring BLA→CeM neurons (paired t-test, t_16_=3.745, \*\**p*=0.0018). n=5 mice, 17 cells. (C) Following food deprivation, within-population photostimulation of BLA→NAc neurons increases current-evoked firing in neighboring BLA→NAc neurons (paired t-test, t_15_=2.217, \**p*=0.0425). n=5 mice, 16 cells. (D) Within-population stimulation of BLA→CeM neurons facilitates firing in neighboring BLA→CeM neurons in slices taken from food deprived mice (paired t-test, t_13_=3.251, \*\**p*=0.0063). n=4 mice, 14 cells. (E) Food deprivation does not impact the interaction between neighboring BLA→NAc neurons in the BLA *(Mann-Whitney U*=115.5, *p*=0.6491). (F) The facilitation of current-evoked firing among BLA→CeM neurons by neighboring BLA→CeM neurons photostimulation is not dependent upon satiety state *(Mann-Whitney U=78.0, p*=0.1085). Dotted rectangles in the example traces shown in (A)-(D) represent the time period of blue light delivery during a current injection, which corresponds to the epochs used to compare firing rates. Bar graphs illustrate mean with SEM. In firing rate difference plots, mean change in firing rate indicated in black circles.

**Figure S6.**
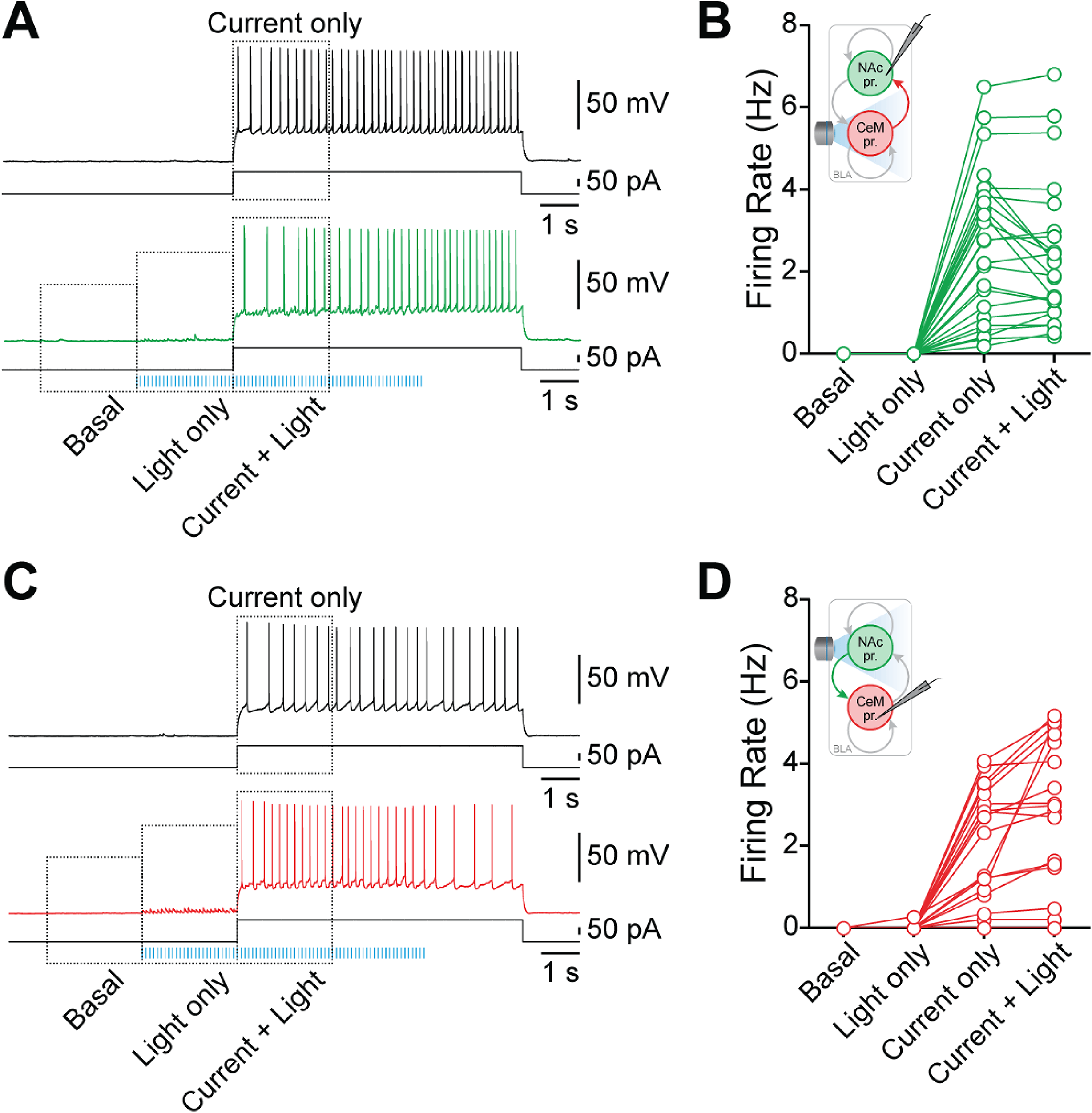
Photostimulation of neither population directly evokes firing in the other (Relates to Main Figure 2) (A) We compared basal firing rate, firing rate in response to photostimulation of the opposing population alone, firing rate in response to current injection only, and firing rate in response to current plus photostimulation. These epochs are illustrated on an example trace from a BLA→NAc neuron during current injection alone (top trace) and during BLA→CeM photostimulation combined with current injection (bottom trace). (B) Firing rate of all BLA→NAc neurons recorded in sated animals in response to BLA→CeM photostimulation during the time periods represented in (A). (C) Example trace from a BLA→CeM neuron during current injection alone (top trace) and during BLA→NAc photostimulation combined with current injection (bottom trace). (D) BLA→CeM firing rate in sated animals in response to BLA→NAc photostimulation during the time periods represented in (C).

**Figure S7.**
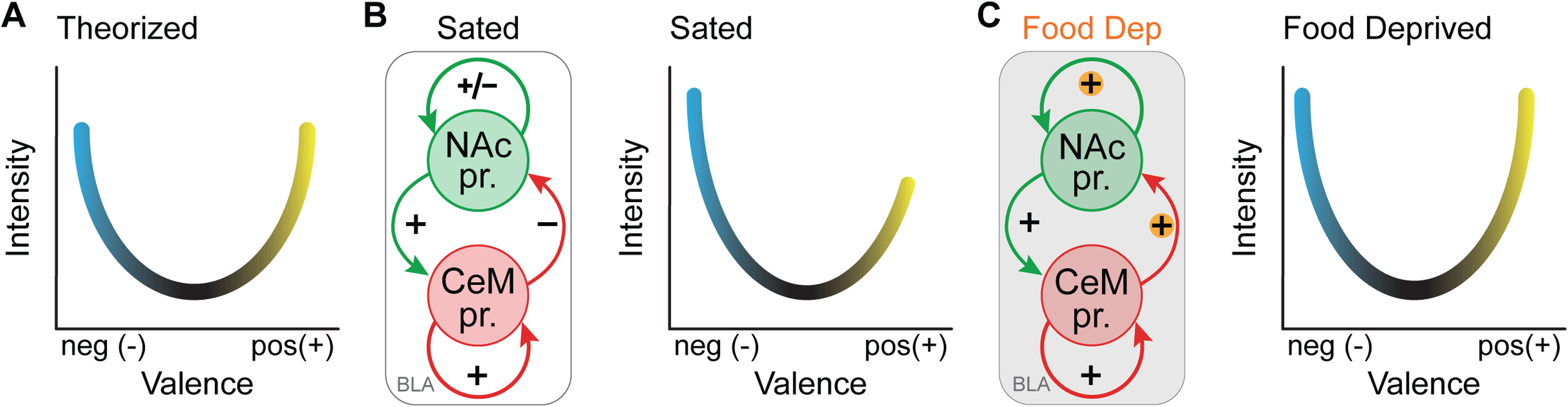
Asymmetric local circuit interactions among valence coding BLA neurons shift depending on satiety state (Relates to Main Figure 2) (A) Based upon our observations that similar synaptic contacts exist between preferentially positive valence encoding BLA→NAc neurons and preferentially negative valence encoding BLA→CeM neurons, we predicted that there would be an equal balance between these two populations, and expected that equal weight would be given to positive and negative valence coding in the BLA. (B) However, we found that in slices taken from sated mice, activity in the BLA→CeM population supersedes activity in the BLA→NAc population. Schematic (left) summarizes these local circuit dynamics between BLA→NAc and BLA→CeM neurons in sated mice. Although both populations are capable of eliciting monosynaptic excitatory and polysynaptic inhibitory responses from each other, BLA→CeM neurons suppress activity in BLA→NAc neurons, whereas BLA→NAc neurons enhance ongoing activity in BLA→CeM neurons. In the schematic presented here, these interactions are represented by + (net excitatory effect), - (net inhibitory effect), and +/- (no net impact on firing rate). This relationship suggests that in sated animals, more weight is given to negative valence coding than positive valence coding (diagram, right). (C) These dynamics are dependent upon satiety state, such that in slices taken from food deprived mice, BLA→CeM neurons no longer inhibit, but rather facilitate activity in BLA→NAc neurons (schematic, left), suggesting that greater weight is given to positive valence coding and reward seeking behaviors in hungry animals (diagram, right).

**Figure S8.**
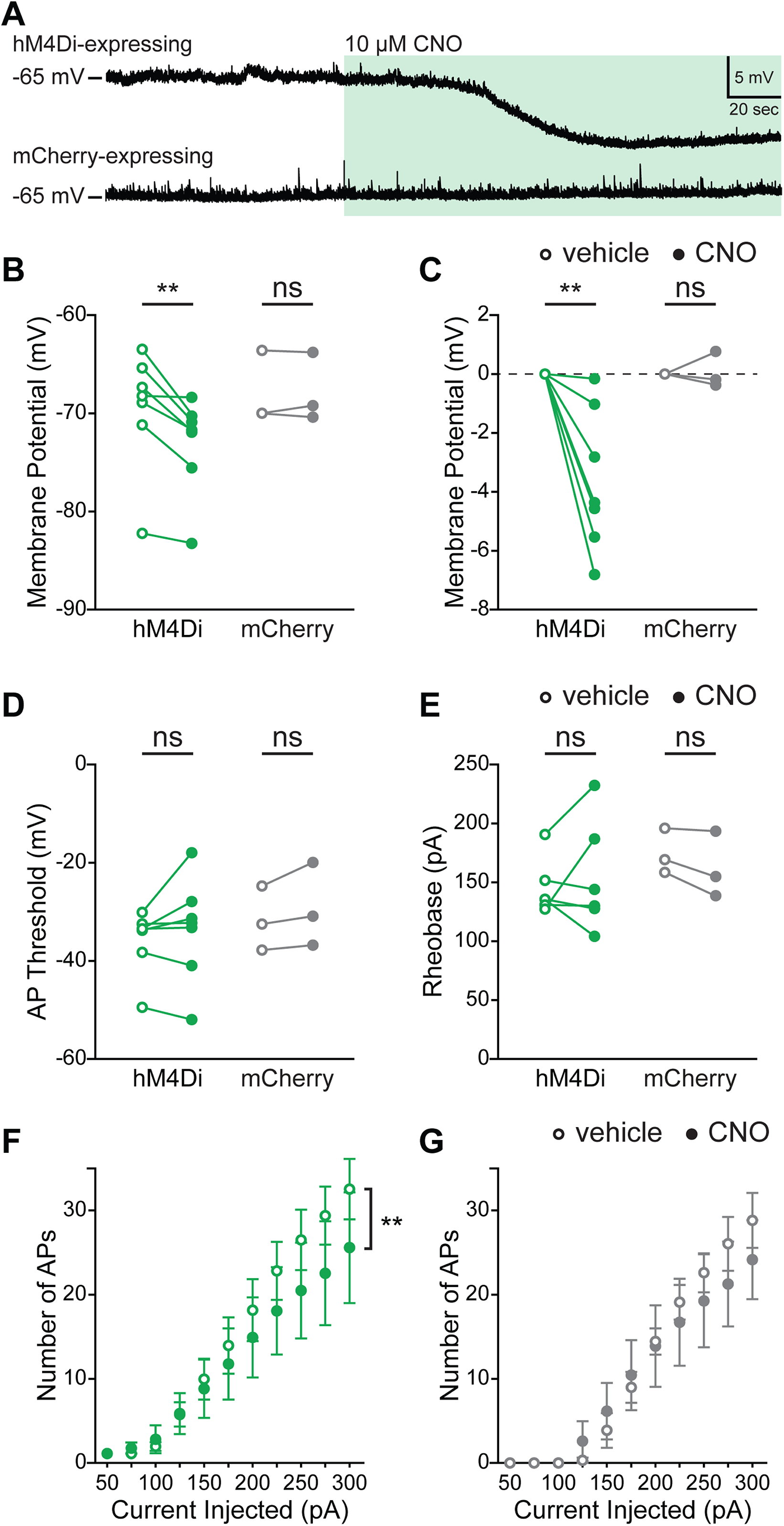
Validation of chemogenetic inhibition of hM4Di-expressing BLA→NAc neurons in acute brain slices (Relates to Main Figure 4) (A) Example traces from whole-cell patch-clamp recordings of hM4Di-expressing (top) and mCherry-expressing (bottom) BLA→NAc neurons in current clamp mode. Bath application of 10 μM CNO resulted in hyperpolarization of the hM4Di expressing neuron, but not the mCherry expressing neuron. (B) Average membrane potential of hM4Di-expressing and mCherry-expressing cells before (open circles) and beginning two minutes after (closed circles) application of 10 μM CNO. CNO significantly reduced membrane potential in hM4Di-expressing cells (paired t-test, t_6_=3.967, \*\**p*=0.0074), but not in the mCherry-expressing controls (paired t-test, t_2_=0.2012, *p*=0.8592). (C) Membrane potential change under bath application of CNO (as in B), shown as relative change from baseline. (D) Action potential threshold, as measured using a 300 pA ramp current test, was unchanged by bath application of CNO in hM4Di- and mCherry-expressing cells (Wilcoxon signed-rank test, *p*=0.4688, Wilcoxon signed-rank test, *p*=0.2500, respectively). (E) The minimum current required to elicit an action potential was also unchanged by bath application of CNO in hM4Di-(Wilcoxon signed-rank test, *p*>0.9999) and mCherry-(Wilcoxon signed-rank test, *p*=0.2500) expressing cells. (F) The relationship between injected current and firing frequency revealed that there was an interaction between drug treatment and current level, indicating that CNO inhibited firing in hM4Di-expressing cells (two way ANOVA, drug*current interaction F_10, 60_=2.444, \**p*=0.0161). (G) There was no impact of CNO bath application on the current/frequency relationship in mCherry-expressing cells (two way ANOVA, drug*current interaction F_10, 20_=0.9005, *p*=0.5499). For all panels, hM4Di: n=7 cells from 3 mice; mCherry: n=3 cells from 2 mice.

**Figure S9.**
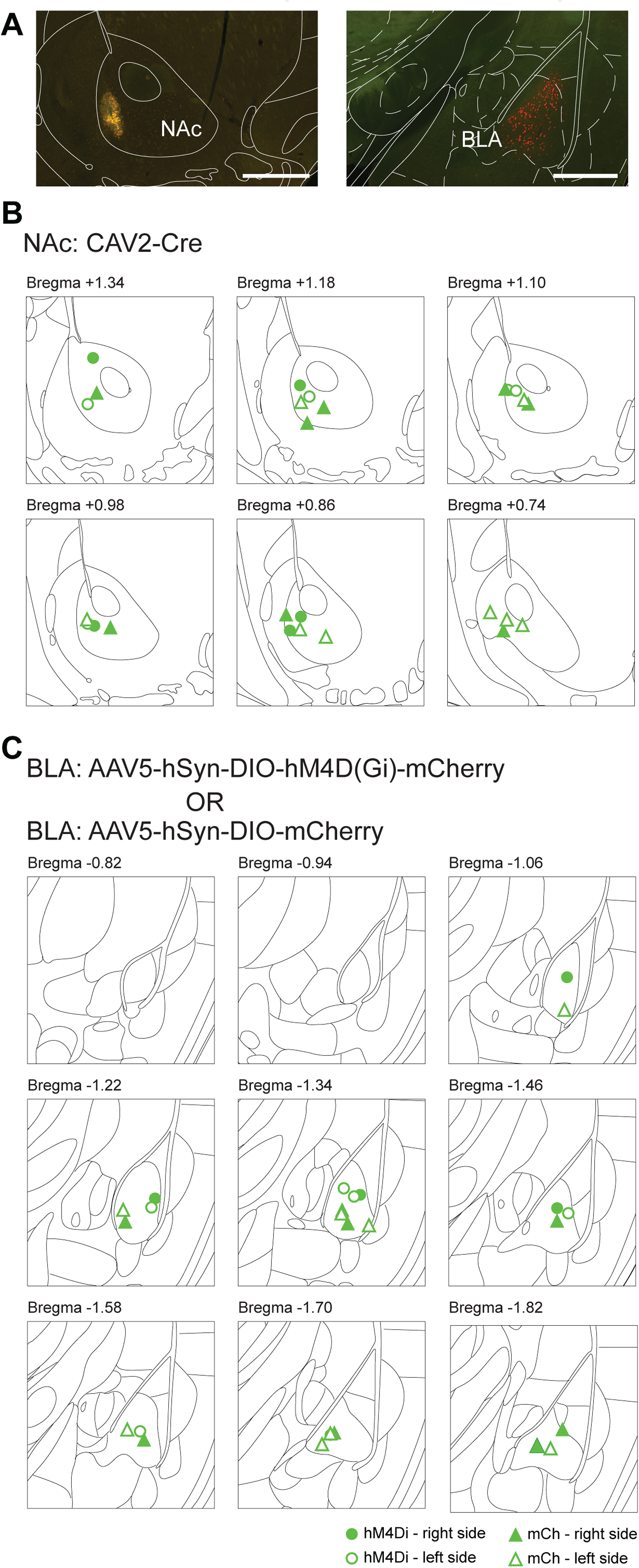
Injection sites for mice used in chemogenetic experiments (Relates to Main Figure 4) (A) Episcope images of CAV2-Cre injection site in the NAc (left), and BLA→NAc neurons expressing hM4Di-mCherry in the BLA (right). Red = mCherry, green = epifluorescence in the eYFP channel, scale bars = 500 μm. (B) Center of CAV2-Cre injection site in the NAc. (C) Center of mCherry fluorescence in the BLA. Closed circles = right hemisphere injection sites from hM4Di mice, open circles = left hemisphere injection sites from hM4Di mice, closed triangles = right hemisphere injection sites from mCherry mice, open triangles = left hemisphere injection sites from mCherry mice.

**Figure S10.**
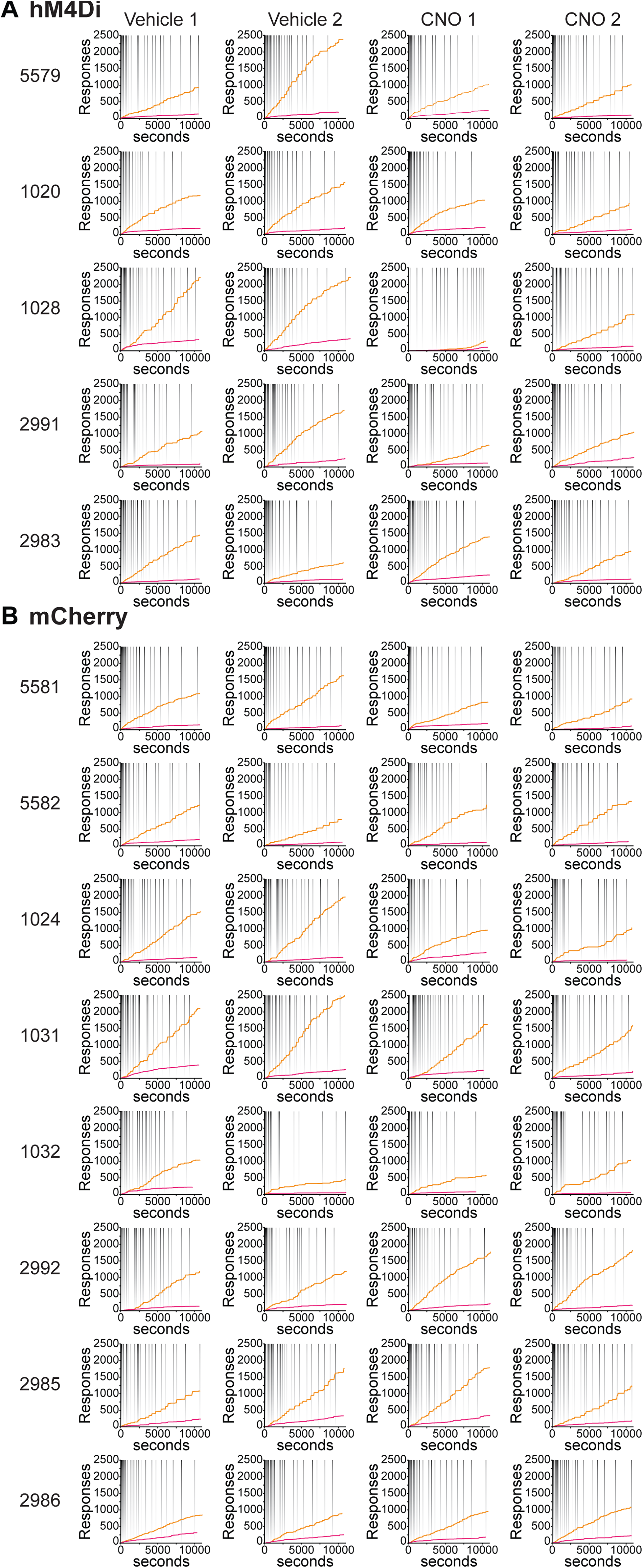
Progressive ratio test day performance for all individual mice (Relates to Main Figure 4) (A) The performance of all animals expressing hM4Di-mCherry in BLA→NAc neurons on the progressive ratio task following administration of vehicle (first two columns) or CNO (second two columns). Each row represents one animal’s behavior. Cumulative nose pokes are indicated in yellow, cumulative reward port entries are indicated in magenta, and reward deliveries are indicated in gray. (B) The performance of all animals in the mCherry control group on the progressive ratio task following administration of vehicle or CNO.

## Author Contributions

Conceptualization, K.M.T. and G.G.C.; Methodology, G.G.C., A.K.S., C-J.C., and K.M.T.; Investigation, G.G.C., A.K.S., C-J.C., G.F.G., C.L.L., A.B., and G.D.M.; Software, C-J.C., A.M.L., P.N., and C.A.L.; Formal Analysis, G.G.C., C-J.C., and A.M.L.; Writing - Original Draft, G.G.C. and K.M.T.; Writing - Review & Editing, G.G.C., A.K.S., C-J.C., C.A.L., G.D.M., C.A.S., and K.M.T.; Visualization, G.G.C., C-J.C., A.B., and K.M.T.; Resources, E.Y.K, C.A.S., C.P.W., A.B., and K.M.T.; Funding Acquisition, G.G.C., A.B., P.N., C.A.L., and K.M.T.; Project Administration, G.G.C. and K.M.T.; Supervision, K.M.T.

## Acknowledgements

We thank the entire Tye laboratory for helpful discussion, and we thank Kaytee Flick for her assistance. We thank E.J. Kremer for providing the CAV2-Cre vector, and the UNC vector core for the AAV 5 vectors. We recognize the generosity of the Genetically-Encoded Neuronal Indicator and Effector (GENIE) program, the Janelia Farm Research Campus, Vivek Jayaraman, Rex A. Kerr, Douglas S. Kim, Loren L. Looger, Karel Svoboda for providing GCaMP6m, and we also acknowledge UPENN for packaging GCaMP6m. We also acknowledge Bryan Roth for providing the inhibitory DREADDs vector to Addgene, from whom we purchased it. K.M.T. is a New York Stem Cell Foundation - Robertson Investigator and McKnight Scholar and this work was supported by funding from the JPB Foundation, PIIF, PNDRF, JFDP, Whitehall Foundation, Klingenstein Foundation, NARSAD, Alfred P. Sloan Foundation, New York Stem Cell Foundation, NIH R01-MH102441-01 (NIMH), and NIH Director’s New Innovator Award DP2-DK-102256-01 (NIDDK). G.G.C. was supported by a NARSAD Young Investigator award through the Brain & Behavior Research Foundation and by the JFDP Postdoctoral Fellowship from the JPB Foundation. A.B. was supported by a fellowship from the Swiss National Science Foundation and NARSAD. P.N. was supported by Singleton, Leventhal, and Whitaker fellowships. C.A.L. was supported by an NSF Graduate Research Fellowship. C.A.S. was supported by a postdoctoral fellowship from the NIH (F32 MH111216, NIMH), and a NARSAD Young Investigator Award (Brain and Behavior Research Foundation).

